# Population Morphology Implies a Common Developmental Blueprint for *Drosophila* Motion Detectors

**DOI:** 10.1101/2025.11.15.688637

**Authors:** Nikolas Drummond, Arthur Zhao, Alexander Borst

## Abstract

T4 and T5 neurons are the first direction-selective neurons in the visual pathway. They are the most numerous cell types in the fly brain (∼ 6000 within each optic lobe) and, as a population, their compact dendritic arbours span the entire visual field. They are classified into four subtypes (a, b, c, and d). Each subtype encodes one of four orthogonal motion directions (up, down, forwards, backwards). Crucially, the dendrites of these neurons are oriented inversely to the functional direction of motion which they encode. This dendritic orientation is what ultimately determines their functional directional encoding.

The development of these neurons is well characterised up to the point of neuropil innervation. However, the full population of these neurons innervate their target neuropil prior to the emergence of directionality within their dendrites. As it stands, development prior to the emergence of dendritic orientations, and the adult oriented dendrite are both well understood, but the key components relating to the emergence of orientation itself are missing.

Recent whole-brain electron microscopy (EM) connectomes of *Drosophila melanogaster* provide an unprecedented level of resolution and completeness when considering the morphology of neurons. Utilising this, we isolate the dendritic arbour of every T4 and T5 neuron within a female adult *Drosophila* brain, made available through FAFB-FlyWire. In doing so we are able to rigorously quantify the morphology of these dendrites in order to understand their similarities and differences.

In doing so we aim to shed light on the origins of dendritic directionality. We reason that either this emerges through a tightly controlled, subtype specific mechanism, or is the result of a subtype agnostic mechanism and external factors. In the former case, we would expect evidence of this in differences between the morphological structure of individual dendrites between T4 and T5, and their subtypes.

Our analysis however reveals a high degree of structural similarity between T4 and T5, and within their subtypes. Particularly, the geometry of branching, section orientation, and tree-graph structure of these dendrites show only minor variability, with no consistent separation between T4 and T5, or their subtypes.

These results indicate that, despite forming in different neuropils, and serving distinct motion directions, T4 and T5 dendrites follow closely aligned morphological patterns. This suggests a shared mechanism of directed outgrowth, as opposed to symmetry breaking emerging through neuron type or subtype specific mechanisms.

**Author summary:** The Dendrites of T4 and T5 neurons orient spatially in a direction inverse to the direction of motion which they encode. The development of these neurons is understood up to the point of neuropil innervation, however they remain unicolumnar. The window between the directed adult dendrite, and their unicolumnar neuropil innervation is not yet understood. In order to shed light on how directionality within these dendrites may emerge, we perform a rigorous quantification of the morphology of all T4 and T5 dendrites within a single female adult *Drosophila* brain using *nm*-resolution reconstructions available from recent electron microscopy.

This allows for the characterisation of neuropil and directionally specific morphological similarities and differences within this population. Although insightful variability is observed at the level of spatial properties of these dendrites, their geometry and topology is highly conserved across T4 and T5 populations, and directions of outgrowth.

From these analyses, we conclude that although these dendrites originate in differing brain regions, with differing input partners, and perform differing, subtype-specific, detection of motion directions, the underlying process of dendritic outgrowth and emergence of orientation is most likely shared across both T4 and T5 and their subtypes.

## Introduction

T4 and T5 neurons are the first motion direction-selective neurons within the *Drosophila* visual system [1, 2]. Integral to the direction-selective function of these neurons is the asymmetric spatial embedding of their dendrites [3, 4]. Our knowledge of the function and structure of these neurons within the adult is well established [1, 2, 5, 6], and their development is understood up to a point [7, 8]. However, how this fundamental asymmetry emerges within their dendrites, and how this impacts structure and function beyond spatial occupancy is not clear. Here we present a comprehensive analysis of the morphology of T4 and T5 dendrites specifically targeted at developing a rich quantitative description in order to evaluate the morphological similarities between these closely related neurons.

These two populations of neurons respond to moving light increments (ON pathway, T4), receiving their inputs within the Medulla, and moving light decrements (OFF pathway, T5), within the Lobula. Both populations of neurons extend their axons to form outputs within the Lobula Plate. The Medulla and Lobula are layered neuropils, with T4 dendrites forming within Medulla layer 10 and T5 dendrites forming in Lobula layer 1 (Fig 1A). Additionally, both neuropils possess a retinotopic mapping through their columnar structure which maps one to one with the lens organisation within the retina [9, 10].

**Fig 1.**
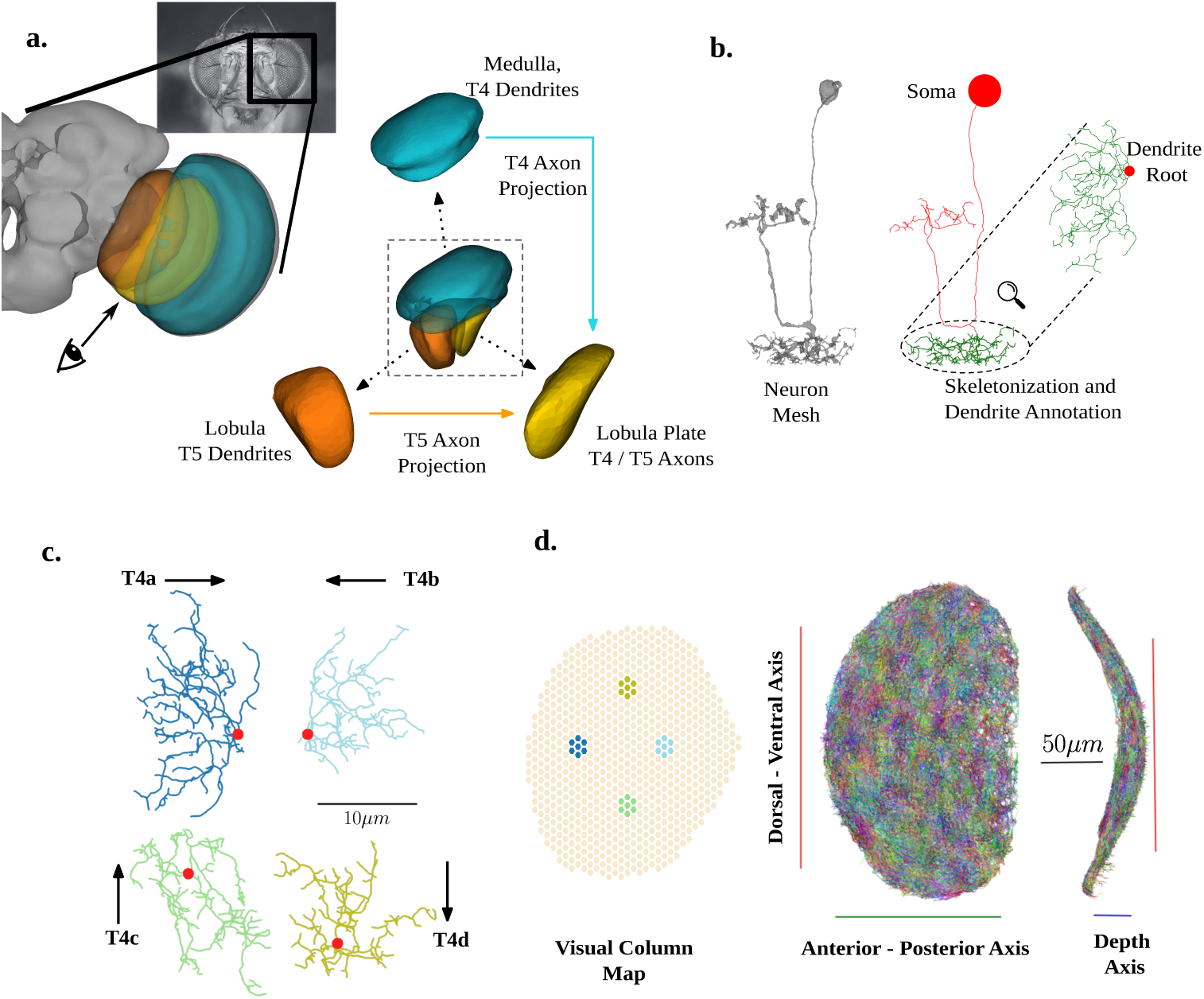
Overview of T4 and T5 dendrites within the *Drosophila* optic lobe. a) (*left*) View of the right optic lobe available in FAFB-FlyWire, with Medulla (Turquoise), Lobula (Orange) and Lobula Plate (Yellow) shown. (*right*) Exploded view of neuropils coloured from perspective illustrated by the eye schematic in *left*. Coloured arrows denote axonal projections for T4 (Medulla to Lobula Plate) and T5 (Lobula to Lobula Plate). b) Illustration of skeletonization and Dendrite annotation procedure from mesh to extracted example dendrite. c) Example extracted T4 dendrites for each subtype. Red points show dendrite root points (first branching point). Arrows show encoded directional tuning of each subtype, inverse to the direction of dendrite outgrowth. d) All extracted T4 dendrites within the right optic lobe in the globally aligned Dorsal-Ventral (DV) / Anterior-Posterior (AP) view, alongside hexagonal grid representation of the *Drosophila* optic lobe columns. Coloured hexagons illustrate Dorsal, Ventral, Anterior, and Posterior example columns used for alignment. Each example dendrite in c matches by colour the T4 dendrite assigned to the centre of each group of coloured hexagons.

Each T4 and T5 neuron acts as a motion detector by integrating visual inputs across adjacent points in visual space [1, 2], with each dendrite spanning across seven columns [2, 4]. Both T4 and T5 neurons form four subtypes (a, b, c, d), which encode the four cardinal directions of body motion (forwards, backwards, downwards, upwards) [3]. Strikingly, each dendrite subtype is physically oriented in a direction inverse to its preferred functional direction. For example, a T4 or T5 ‘a’ subtype will grow along the anterior-posterior axis, from posterior to anterior, but encode motion anterior to posterior (Fig 1C). This orientation is determined by the structure of the compound eye, and is dominant within the central region of the eye [10]. Additionally, evidence in favour of a further functional subtype division along the horizontal axis (a and b subtypes) is given in [11].

Considering the relevant development of T4 and T5 neurons, the four optic lobe neuropils within *Drosophila* (the Lamina, Medulla, Lobula, and Lobula Plate) develop during the larval and early pupal stages from two neuroepithelial domains, the inner and outer proliferation centres. The T4 and T5 neurons are produced by progenitors situated within the proximal inner proliferation centre [7, 12] and, through a series of binary switches a single progenitor will form into a complement of four neurons. Specifically, two T4 and two T5 along either the horizontal (forwards and backwards motion, a and b subtypes) or vertical (upwards and downwards motion, c and d subtypes). During this developmental cascade, the final type division separates T4 and T5 neurons of the same subtype, for example, T4a and T5a [7]. Both T4 and T5 dendrites innervate their target neuropils in a spatio-temporal wave, forming innervations along the anterior-posterior axis over the same time window. Additionally, neurons from a shared progenitor form in the same region of both the Medulla and Lobula through this wave [7].

By 36 hours post puparium formation (APF) the subtype specific axonal projection into distinct layers of the Lobular Plate is established, and remains stable throughout the remainder of the developmental period [8]. At 36 hours APF the dendrites of both T4 and T5 neurons have fully innervated their target neuropils (the Medulla and Lobula respectively), however are, at this stage, uni-columnar, and show no direction selectivity [8]. As such, going from Larval to early pupal stages, up to 36 hours APF, all T4 and T5 dendrites are situated within their target location, and have their specific subtype identity, but do not show directional selectivity within their dendrites.

Between 36 and 72 hours APF, directional selectivity within these dendrites emerges simultaneously at an equivalent rate across all T4 and T5 and their subtypes before undergoing a a period of pruning between 72 hours APF and full maturity as a adult [8]. During this critical developmental period (36-72 hours APF) the mechanism behind how the directionality of these dendrites emerges remains a mystery. The first step towards resolving this knowledge gap requires the accurate characterisation of the dendritic arborisation patterns, focusing on which properties are conserved across differently oriented subtypes, and which are orientation specific.

This is a particularly pressing concern within the T4 and T5 neurons, as their directional encoding, and by extension their entire role in computation, is determined by the spatial embedding of their dendrites [1, 2, 5, 6]. In order to move our understanding of the emergence of directionality within these dendrites forwards, we turn to a fully reconstructed connectome of a female adult *Drosophila* brain (FAFB-FlyWire) [9, 13–16]. This provides us with *nm* resolution reconstructions of all T4 and T5 adult neuron morphologies in both hemispheres, *n ∼* 11000 neurons. By isolating their dendrites we are able to perform in-depth analysis of their morphology, isolating the features of their spatial embedding, geometry, and topology to compare similarities and differences.

From these analyses, we envision two outcomes which offer differing interpretations of the emergence of directional selectivity. First, it is possible that symmetry breaking is a result of subtype specific mechanisms which directly inform the structure of each dendrite. As such, we would predict subtype specific variability within the geometry and topology of each dendrite, alongside differences in their spatial embedding. This would indicate a tightly controlled, subtype specific, mechanism of dendrite outgrowth during the period between 36 and 72 hours APF. On the other hand, symmetry breaking could be the result of subtype agnostic mechanisms. In such a case, we would expect homogeneity within the geometric and topological structure of each dendrite subtype, with observable differences being primarily in the nature of the spatial embedding of each dendrite, which is already known in relation to these dendrites asymmetry.

When considering the spatial embedding of these dendrites, a number of small type and subtype specific differences are observed beyond their characteristic orientation. Strikingly however, in the latter two sections of our analysis we find no systematic differences between T4 and T5, or their subtypes. As fine-grained, local, structure does not differ between these dendrites, this leads us to conclude that functional directionality is determined solely by spatial embedding, and not through morphological differences between types and subtypes. Second, this similarity is indicative of the hypothesis that the manner of dendritic outgrowth is shared across T4 and T5 dendrites, as well as their subtypes, and symmetry breaking is a result of factors independent of the dendrite outgrowth itself.

## Results

### Note on Statistical Analysis and Outcomes

Due to the large *n* available within this dataset, standard practices relying on reporting *p*-values are not suitable here (See Materials and Methods, Statistical Analysis for further discussion). As such, we focus on reporting effect sizes in order to emphasise the magnitude of differences over statistical significance alone [17, 18]. For main effects and interactions we use partial eta squared (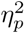), In order to define a meaningfully large effect size we use guidelines from [19]. These are as follows: “small” effect size as ≥ 0.01, a “medium” effect size as ≥ 0.06, and a “large” effect size as ≥ 0.14. We use these as a cut off, and require that a reported meaningfully large effect size, and the confidence interval around it, is typically above the “small” cut-off.

Similarly, for *post hoc* comparisons, we report Cohen*′*s*_d_* (”small”: ≥ ±0.2, “medium”: ≥ ±0.5, “large”: ≥ ±0.8) with the same criteria.

### Dendrite Spatial Embedding

We first turn to understanding the spatial structure and embedding of T4 and T5 dendrites in order to understand differences in the manner in which these dendrites occupy their densely overlapping environment. Here we focus on the shape and volume of the spanned region of T4 and T5 dendrites, the manner in which these dendrites innervate across the depth axis of their respective neuropil layers, and the spatial relationships between the first branching point of different dendrites.

#### Dendrite Shape

T4 and T5 dendrites are often characterised by their asymmetrical orientation [3], whereby the direction of dendrite outgrowth is inverse to the direction of functionally encoded motion (Fig 1C). In order to visualise the occupied space of dendrites by subtypes we align individual dendrite in a manner which allows us to overlay them within subtypes. For each dendrite we specify its internal axis using Eigendecomposition of the underlying point cloud representation of each dendrite. As the covariance matrix of 3-dimensional coordinates is a square symmetric matrix, the eigenvectors are orthogonal to each other, and relate directly to vectors in space for each dendrite - their internal axis. We can then order axis based on the magnitude of their corresponding eigenvalues. We term the axis corresponding to the largest eigenvalue as the ‘longest’ internal axis, as this is the axis which explains the greatest variance within the data. As these data are points in 3-dimensional space this will be the axis of longest extent spatially. We orient each ensuring that the longest axis aligns with the y-axis basis (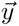 = [0, 1, 0]), and scale by the mean standard deviation along the y-axis within each subtype (See Materials and Methods, Alignment and Scaling). By then centring individual dendrites on the root of their dendritic branching (referred to as the dendrite root point) we can overlay all dendrites within subtypes (Fig 2A). Additionally, as the spatial coordinates are scaled to one standard deviation along the y-axis within each population, we can overlay both hemispheres within the same space. This shows us a clear elliptical spatial occupancy shared across all dendrite subtypes. Having centred dendrite subtype populations on their dendrite root point, we also observe clear offsets from the centre of each elliptical shape for each subtype (Fig 2A; Red points), with the exception of T5a subtypes, where the dendrite root point appears more centrally positioned. Additionally, in comparison to T4 c and d subtypes, the dendrite root point of T5 c and d subtypes appears additionally offset towards the anterior. When this alignment is performed separately within each hemispheres dataset (Fig S1), the same offset patterns in root positions are observed.

**Fig 2.**
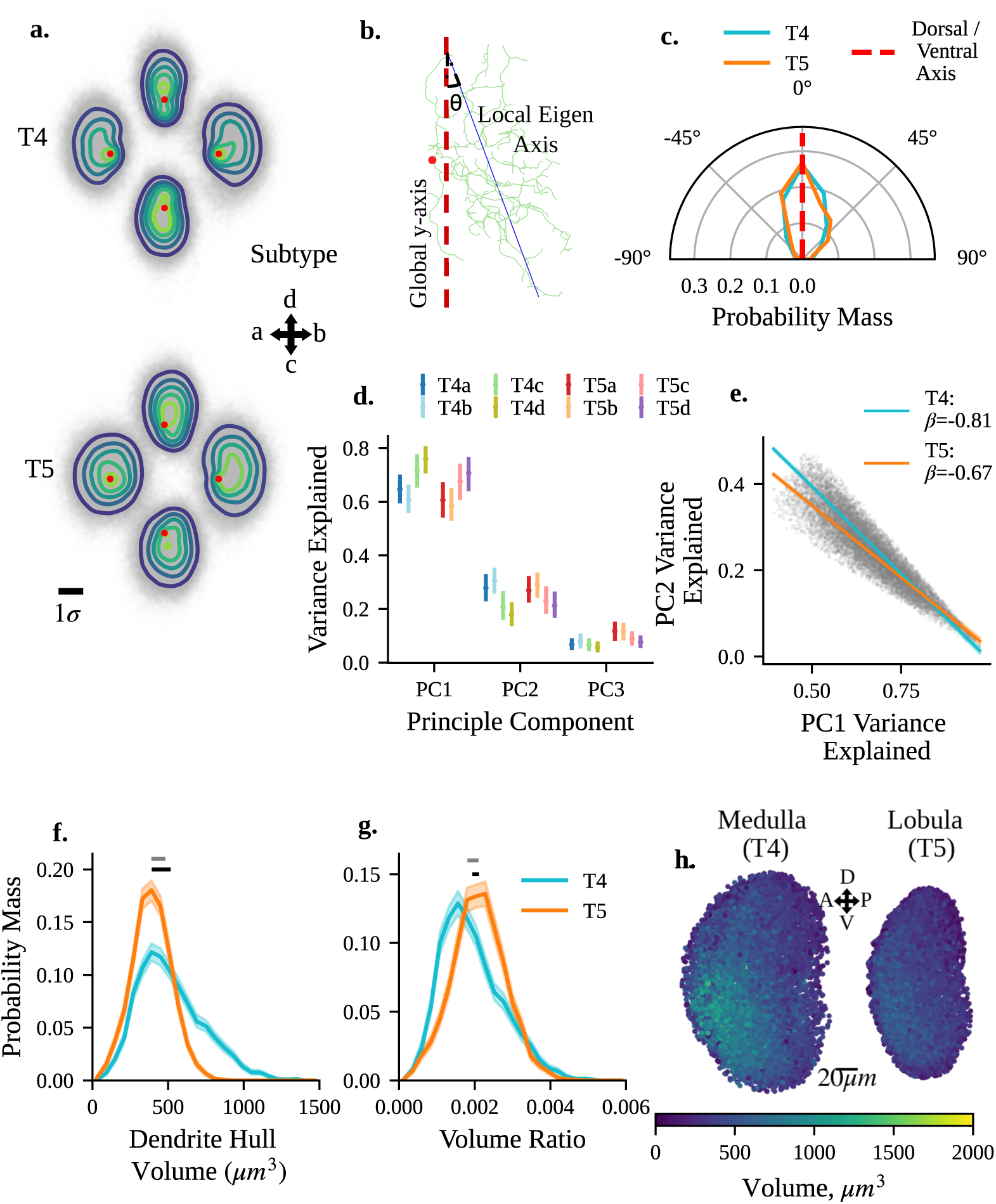
Spatial Embedding and Volume of T4 and T5 Dendrites. a) Scaled and locally aligned subtype specific dendrite populations, pooled over hemispheres. Each individual dendrite is aligned to its internal axis and scaled within each type specific population (T4 and T5) to one standard deviation of the population dispersion along the longest axis. Individual dendrites are centred on the root of the dendritic tree (red points). KDE’s show point density for each subtype. b) Schematic illustrating angular deviation between a dendrites longest internal axis (blue line) and the global Dorsal-Ventral (DV) axis (red dashed line). Red point shows the dendrite root. c) Distribution of angles between DV axis and dendrites longest internal axis for T4 and T5 dendrites d) Principal component analysis variance explained along each principle component (PC) for each subtype, illustrating dispersion of points along each internal axis per dendrite. e) Relationship between variance explained between first and second principle components for T4 and T5 dendrites. f) Distribution of raw convex hull volumes for T4 and T5 dendrites. Black horizontal lines denotes difference in means. Gray horizontal lines denote difference in medians. g) Distribution of convex hull volumes scaled by Medulla layer 10 volume (T4) and Lobula layer 1 volume (T5). Black horizontal lines denotes difference in means. Gray horizontal lines denote difference in medians. h) Spatial distribution of T4 and T5 convex hull volumes within their respective neuropil (T4, Medulla, left; T5, Lobula, right) on the globally aligned DV / AP plane collapsed together for both hemispheres.

Given these dendrites’ characteristic asymmetry, it is unclear from this alone if the longest internal axis, and the extension of these dendrites in space, is shared along either the dorsal-ventral (DV) or anterior-posterior (AP) global axis across all subtypes, or differs between horizontally oriented (a, b) and vertically oriented (c, d) subtypes. In order to investigate this we measure the angle between each dendrites’ longest internal axis and the global DV axis (Fig 2B). We find that the longest internal axis of all neuron types and subtypes in both hemispheres is aligned well with the global DV axis (Fig 2B). There is no meaningfully large difference between types, subtypes, or hemispheres (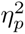 ≤ 0.01). This elongation is in line with the observed elongation of columns along the DV axis in both Medulla layer 10 and Lobula layer 1 [4].

We turn next to understand if these dendrites differ in their spread along the DV, AP, and depth axis. We extend the Eigendecomposition utilised for alignment to principal component analysis (PCA). As in standard PCA, the variance explained by each principal component is given by dividing each eigenvalue by the sum of eigenvalues. This tells us the amount of dispersion within the point cloud representation of dendrites along each of its internal axes. As we have shown already that the longest internal axis aligns well with the global DV axis, it follows that the central internal axis aligns with the AP axis, and the shortest internal axis with the depth axis globally.

The variance explained by PC1 we find no meaningfully large difference between neuron types or hemispheres 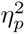 ≤ 0.01. Between subtypes we find a medium effect size (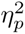 = 0.086, CI : [0.075, 0.099]). *Post hoc* pairwise comparisons find all comparisons are meaningfully large (Cohen*′*s*_d_* > 0.2). Considering the mean differences between subtypes, there is a clear structuring in this result. The variance explained along the first axis, which translates to the DV axis, is ordered in a subtype-specific manner following *d* > *c* > *b* > *a*.

Similarly, for variance explained by PC2 we again find no meaningfully large difference between neuron types or hemispheres (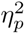 ≤ 0.01). We again observe a medium effect size between subtypes (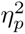 = 0.088, CI : [0.078, 0.101]). *Post hoc* comparisons are all have a meaningfully large effect size, with the exception of a vs b subtypes, where the tail of the confidence interval falls bellow the small effect size threshold (Cohen*′*s*_d_* = −0.243, CI : [−0.192, −0.296]. Interestingly however, the previously observed ordering between subtype means is reversed (*d < c < a < b*).

Within PC3 we find a slightly different result. Here, we observe a small effect of neuron type (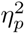 = 0.033, CI : [0.027, 0.041]). *post hoc* comparisons show that T4 ¡ T5 with a medium effect size (Cohen*′*s*_d_* = −0.754, CI : [−0.719, −0.790]). This difference may be a result of the two different depths of the respective neuropil layers being innervated by T4 and T5. No meaningfully large differences are observed between neuron subtypes or hemispheres.

Given the orderings observed in variance explained by PC1 and PC2, we fit an ANCOVA model for PC2 with PC1 as a covariate (Fig 2E). Here we find a large main effect indicating PC1 as a strong predictor of PC2 (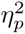 = 0.184, CI : [0.154, 0.223]), with a slope of *β* = −0.806. Here, no meaningfully large differences are observed between neuron types, subtypes, hemispheres, or interactions.

Together, these results tell us that all T4 and T5 dendrites elongate primarily along the DV axis in both the Medulla and Lobula. However, the extent of this elongation is subtype-specific, although in the same manner between T4 and T5. Interestingly, the observed negative correlation between the variance explained along PC1 and PC2, which relate to the DV and AP axis, indicates that the greater the extent of outgrowth along the DV axis, the lesser the extent along the AP axis in a proportional manner.

#### Dendrite Volume

Having established the shape of T4 and T5 dendrites we next turn to investigate the spanned volume of each dendrite. This is done by fitting a convex hull to the each individual dendrite morphology (Fig 2F). Here we find a small main effect of neuron type (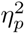 = 0.022, CI : [0.018, 0.028]). *Post hoc* comparisons indicate a medium sized effect indicating that mean convex hull volume is greater in T4 than in T5 (Cohen*′*s*_d_* = 0.686, CI : [0.652, 0.712], Fig 2F). No meaningfully large effects are observed between neuron subtypes, hemispheres, or interactions.

However, the Medulla is notably larger than the Lobula. In order to account for this difference, we scale dendrite convex hull volumes by the volume of hemisphere specific surface meshes for Medulla layer 10 for T4 and Lobula layer 1 for T5 (see Materials and Methods, Surface Fitting, Volume, and Depth Measurement). Once doing so, all neuron type, subtype, hemisphere, and interaction effect are no longer meaningfully large 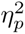 ≤ 0.01 (Fig 2G). This indicates that the mean proportional volume is the same across all types and subtypes, as would be expected from each dendrite spanning seven adjacent columns in space. Additionally, we note that the peak of proportional volumes observed (≈ [0.001, 0.002]) is within the range expected by partitioning the neuropil layer volume into 700 equal sized regions (1/700 = 0.0014).

Additionally, the use of convex hull volume around each dendrite is only one measure of the size of an individual dendrites. We additionally consider the total cable length of each dendrite, as well as the number of sections between the dendrite root point, branching nodes, and terminal (leaf) nodes as additional metrics of size (S2 Fig). It is interesting to see that each of these size-based metrics is highly correlated, illustrating the consistent relationship between size based metrics within these dendrites.

We note however that the distribution of T4 volumes is notably more heavy tailed than that of T5 (Figs 2F and 2G, S2). When we look at the volume of T4 and T5 dendrites plotted spatially, we note a clear structure in T4 not present in T5 (Fig 2H; T4: left, T5: right). The tail of larger T4 dendrites clearly cluster in the anterior-ventral region of Medulla layer 10, a spatial organisation which is not clearly visible in T5 dendrites within Lobula layer 1. This observed pattern is mirrored in other size-based morphometrics used, including total dendrite cable length, and the number of sections within each dendrite (the number of edges in the underlying tree-graph, S2 Fig).

This spatial distribution of T4 dendrite size leads to two distinct possibilities. Either the size of columns within the Medulla is non-uniform, with larger columns within the anterior-ventral region, or column sizes are uniform, and individual T4 dendrites within the anterior-ventral region integrate visual inputs across a larger region of the visual field. In order to investigate this we isolate the portion of the surface mesh of each Mi1 neuron within Medulla layer 10 and fit a cylinder, giving us the spatial extent of each column (S1 Text, Materials and Methods). We find that the mean inter-column distance, length, and radius of each column within Medulla layer 10 is uniform however (S3 Fig). This is then evidence in favour of the latter argument; that T4 dendrites within the anterior-ventral region integrate their inputs over a greater proportion of the visual field. That larger dendrites are observed within the region of space which correlates with larger lenses on the retina seems unlikely to be coincidental, however further work investigating functional tuning will be needed to understand the implication of the larger dendrite sizes observed here.

We have shown here that both T4 and T5 dendrites occupy the same mean proportional volume within their differing neuropils, in line with previous work that has shown that T4 and T5 dendrites span across approximately seven visual columns in both the Medulla and Lobula [2, 4]. For T4 dendrites, however, we see a heavy tailed distribution of dendrite size not present in T5, and a spatial organisation favouring larger regions spanned by the dendritic arbour in the anterior-ventral region of Medulla layer 10 which is not apparent in Lobula layer 1. Given the uniformity of column sizes, this would seemingly indicate that T4 dendrites within the anterior-ventral region span across marginally more than 7 columns, however how this may impact function is unclear from these data alone.

Finally, the highly correlated relationships observed between convex hull volumes, total cable length, and total dendrite sections, are indicative of coordinated proportional growth. As dendrites expand the volume of their spatial occupancy, they increase their cable length and branching complexity proportionally.

#### Dendrite Depth

We have focused thus far on the manner by which T4 and T5 dendrites fill space along the DV and AP axes. We turn next to understanding how individual dendrites occupy space along the remaining depth axis. In order to do so, we calculate the normalized neuropil layer depth of dendrite nodes (Fig 3A; Materials and Methods, Surface Fitting, Volume, and Depth Measurement). When globally aligning each population of dendrites, we use a shared origin determined by the centre of mass of a sphere fitted to each population. Given the surface mesh of each neuropil layer, we can determine the inner and outer surface of each mesh based on the dot product of face normals within the mesh to a ray cast from this shared origin, with values less than −0.8 giving faces on the inner surface, and greater than 0.8 providing the outer surface. In the case of the Medulla, the outer surface corresponds to the boundary between Medulla layer 10 and Medulla layer 9, and the inner surface the outer boundary of the Medulla and the surface through which the T4 dendrites innervate. This is inverted within T5, where the inner surface corresponds to the boundary between Lobula layer 1 and Lobula layer 2, and the outer surface to the boundary of the Lobula, and the surface through which T5 dendrites innervate the neuropil. This inversion is why depth is inverted within T5. As a result of the mesh fitting procedure, an offset along the depth axis is introduced between the two hemispheres (See Materials and Methods, Surface Fitting, Volume and Depth Measurement). As a result, analysis on the left and right hemisphere datasets are treated independently here. This is due to the fact that this offset dominates the results showing a misleading difference in hemispheres, however the observations of interest remain consistent between the two hemispheres.

**Fig 3.**
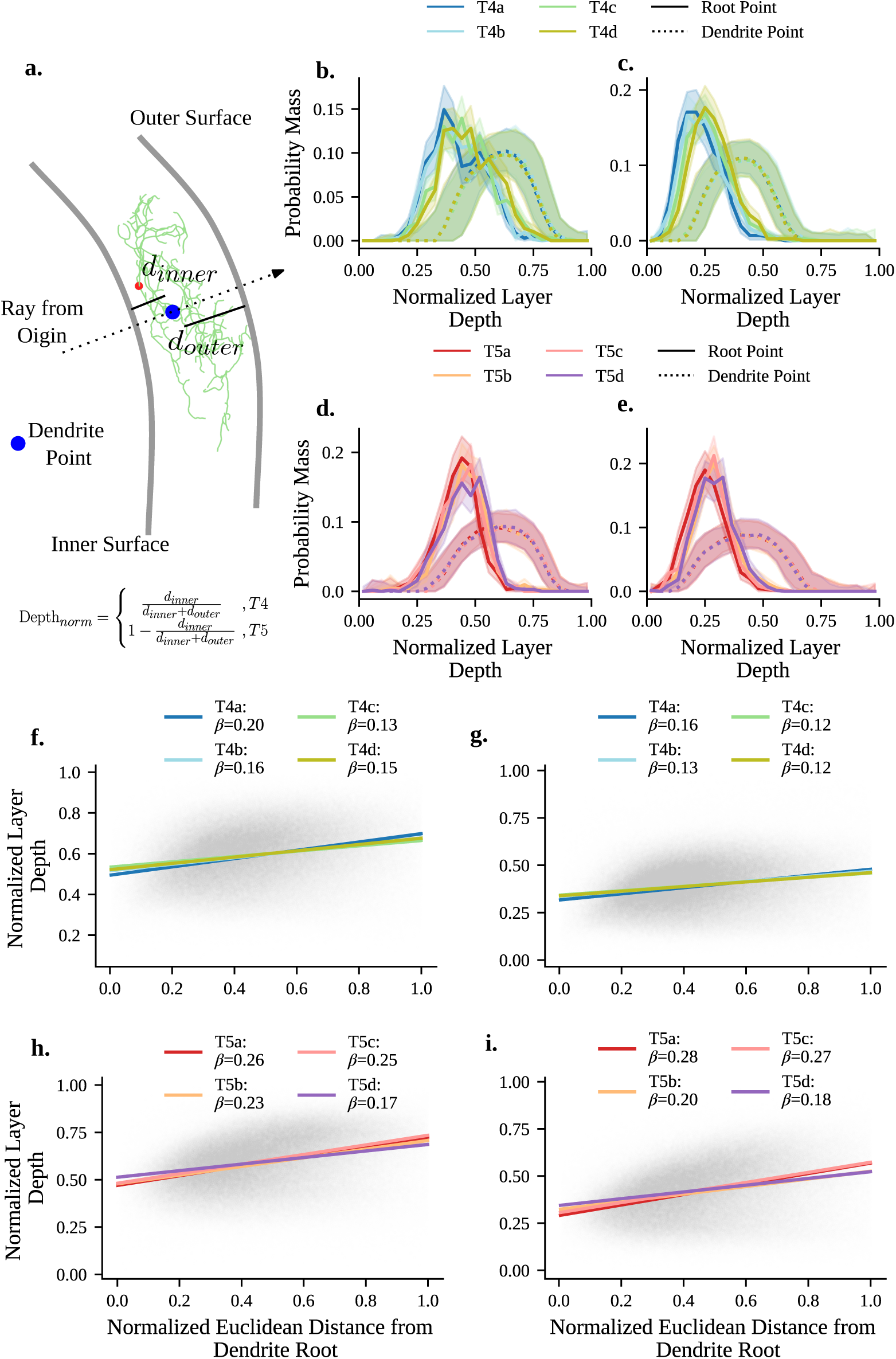
Dendrite Neuropil Layer Specific Depth. a) Schematic illustrating Neuropil specific depth calculation within T4 and T5. Layer surface meshes are generated using all dendrite points, and inner and outer faces estimated using the dot product of a ray from the coordinate origin and the normal of each surface face (See Materials and Methods, Surface Fitting and Volume). The inner surface within T4 dendrites corresponds to the boundary of the Medulla, and the surface through which T4 dendrites innervate their neuropil. The outer surface is the boundary between Medulla layer 10 and Medulla layer 9. Within T5 dendrites, this is inverted, so the inner surface represents the boundary between Lobula layer 1 and Lobula layer 2, and the outer surface the boundary of the Lobula through which T5 dendrites innervate their neuropil. As T5 dendrites innervate from the side of the outer surface, and T4 from the inner, T5 normalized depths are inverted. b) Distribution of T4 dendrite point depths within the left hemisphere. Solid lines indicate dendrite root points, with a bootstrapped confidence interval around the probability mass function. Dashed lines represent the distribution of non-root points within dendrites, shaded region show asymmetric MAD within the probability mass function across individuals. c - e) As in b, for T4 dendrites within the right hemisphere (c), T5 dendrites within the left hemisphere (d), and T5 dendrites within the right hemisphere (e). f) Multiple Regression showing subtype specific slopes within T4 dendrites within the left hemisphere illustrating the relationship between normalized euclidean distance from root (normalised by the maximum per dendrite) and normalized point depth within Medulla layer 10. Root and furthest points within each dendrite (*x* = 0 and *x* = 1) are excluded. g-i) As in f, but for T4 dendrites within the right hemisphere (g), T5 dendrites within the left hemisphere (h), and T5 dendrites within the right hemisphere (i).

We first investigate the depth of the dendrite root point of individual dendrites (Dendrite root, Fig 3B to 3E; solid lines). Within the right hemisphere, we observe a small main effect of neuron subtype (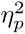 = 0.021, CI : [0.016, 0.026]). This same small main effect is observed within the left hemisphere (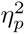 = 0.014, CI : [0.011, 0.021]). *Post hoc* comparisons of both hemispheres reveal this is caused by the ‘a’ dendrite subtype root positions being positioned marginally closer to the neuropil layer surface in comparison to other subtypes. Within the right hemisphere, *post hoc* comparisons show the ‘a’ subtype is closer to the neuropil surface in comparison to all other subtypes with a small to medium effect size (−0.2 > Cohen*′*s*_d_* < −0.8). Within the left hemisphere, this difference is only observed between the ‘a’ and ‘d’ subtypes with a small effect size (Cohen*′*s*_d_* = −0.347, CI : [−0.276, −0.377]). We note that within the left hemisphere, the depth of T4 dendrite root points is significantly more varied and noisy than in the right hemisphere, or T5 dendrites (Fig 3B). This is unlikely to be a result of the dendrite extraction procedure as all dendrite root points are reviewed manually in a random order across the whole dataset. It may be a result of varying reconstruction quality of T4 neurons within the left hemisphere of the FAFB-FlyWire dataset, but it is unclear if this is the case or this is a biological difference at this stage.

When considering the depth of non-dendrite root points we find no differences between type, subtype, or interaction (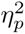 ≤ 0.01) within both hemispheres (Fig 3B to 3E; dashed lines). This indicates that individual dendrites, excluding the start of dendritic branching, occupy the same depth within the neuropil layer regardless of type and subtype, or direction of innervation within the neuropil layer itself.

We next consider the manner by which T4 and T5 dendrites project across their respective neuropil layer. The previously mentioned offset introduced within the layer meshes is most obvious here in the shift along the y-axis (Fig 3F to 3I). As both Medulla layer 10 and Lobula layer 1 are well approximated by a curved surface, two possibilities exist. Either, dendrites could follow this curvature as they fill space, or they could project across this curvature. In the latter case, we would expect each dendrite to exist primarily on a 2D plane, and fill space at an angle across the neuropil layer curvature. Given the proximity of the dendrite root point to their respective neuropil layer surface we would expect dendrite points further from this point to be closer to the opposing layer surface. We include the normalised euclidean distance from the dendrite root point, normalised by the maximum within each dendrite (referred to a root distance), as a covariate predictor for normalised depth within each hemisphere. Within both hemispheres, we find a small main effect of root distance as a covariate predictor of neuropil depth. It is noted that within the right hemisphere, the confidence interval around this effect size drops marginally below our small effect size cut off (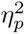 = 0.010, CI : [0.009, 0.010]), with a slope of *β* = 0.183. Within the left hemisphere, this effect is more pronounced (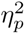 = 0.012, CI : [0.012, 0.013]) and a slope of *β* = 0.219. With no meaningful effects of neuron type or subtype, this indicates that all T4 and T5 dendrites project across the neuropil layer at a slight angle (Fig 3F to 3I), with increasing depth in the neuropil layer away from the dendrite root. We note however that these data are exceptionally noisy and caution is taken with interpretation.

Taken together our analysis shows that the depth at which dendrite subtypes begin arborising may differ by subtype, more noticeably for T4 than T5 subtypes. The depth of the dendritic arbour itself, excluding the dendritic root point, does not differ between types and subtypes. Finally, we find that these dendrites project across their respective neuropils, rather than growing along the curved space.

#### Dendrite Initial Positions

A single column within either Medulla layer 10 or Lobula layer 1 contains the initial innervation of a full complement of all four subtypes of T4 or T5 dendrites [1]. With the absence of neuropil specific spatial column information, we investigate the pairwise nearest-neighbour relationships between dendrite root points. This informs us about any possible spatial structure in the initial starting positions of dendritic arborisation. We find both the subtype-specific nearest-neighbour pairings for dendrite root points, and additionally the global subtype-agnostic unique pair assignment which minimises the total distance between pairs of dendrite root points (Fig 4A; Materials and Methods, Dendrite Root Nearest Neighbour Matching). Alongside subtype labels, we additionally group data into “same” (e.g a→a), “opposite” (e.g a→b), or “orthogonal” (e.g a→c and a→d) groups.

**Fig 4.**
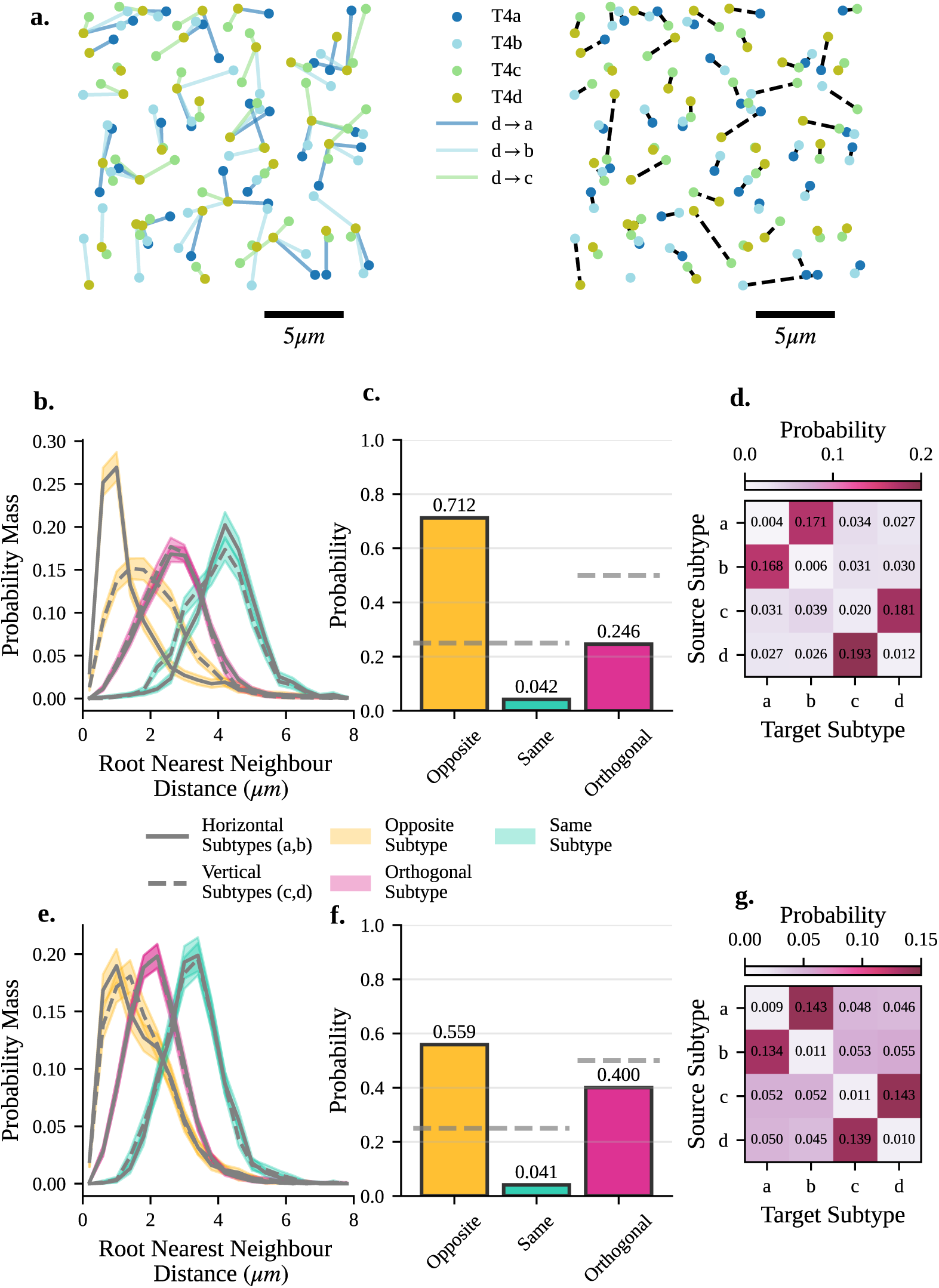
Dendrite root nearest neighbour relationships. a) Illustration of nearest neighbour calculations from a 20*µm* section of dendrite root positions within the central medulla. (*left*) shows subtype specific nearest neighbours from T4d dendrites to T4 a,b, and c subtypes (T4d→T4d are omitted from the schematic for simplicity. (*right*) Global subtype agnostic nearest neighbour calculations determined by maximum cardinality matching in order to find the unique set of nearest neighbour pairs which minimises the global distance between pairs of dendrite root points. b) Nearest neighbour distance of dendrite root points within T4 dendrites, using matching as in (a, left), grouped into horizontal (a, b subtypes, solid lines) and vertical (c, d subtypes, dashed lines). “Opposite” distributions denote opposingly oriented subtypes (eg, a→b). “Same” subtypes denote same subtype combinations (eg, a→a). “Orthogonal” denotes orthogonally oriented subtypes (eg, a c and a d). c) Probability of the nearest neighbour being of each of the “Opposite”, “Same”, and “Orthogonal”, inferred using nearest neighbour matching derived from (a, right) within T4 dendrites. Dashed lines denote approximate chance probability for each group (0.25 for “Opposite” and “Same” groups, 0.5 for “Orthogonal” groups). d) Subtype specific nearest neighbour probabilities inferred from nearest neighbour matching derived from (a, right) within T4 dendrites. e-g) as in b-d but for T5 dendrites.

When looking at the subtype specific distribution of nearest-neighbours, we find that in all cases the “opposite” subtype is the closest nearest neighbour for any given dendrite’s root point, and the “same” subtype is always the furthest, with “orthogonal” subtypes occupying the middle distance, with no separation between which of the “orthogonal” subtypes is closer (Fig 4B and 4E). Additionally, we observe that within T4 but not T5 dendrites, the horizontal (a,b; Fig 4B, solid lines) and vertical (c,d; Fig 4B dashed lines) subtype distributions differ. The vertical subtypes show a greater distance between their nearest neighbours in relation to the horizontal, which is not the case in T5 dendrites (Fig 4E). One possibility is that this difference within T4 is a result of differing neuropil layer depths of the dendrite root points between subtypes. In order to asses this, we repeat this analysis on dendrite root points projected onto the surface of a fitted sphere, fit individually to each neuron type population within each hemisphere (See Supplementary S1 Text, Materials and Methods, Spherical Surface Projection). In doing so, all points are projected to the same depth, removing this as a possible confound. We find no difference in our observed results (S4 Fig).

Based on the subtype agnostic nearest-neighbour pairing, we can additionally calculate the probability of a given nearest neighbour being of the “Opposite”, “Same” or “Orthogonal” classification (Fig 4C and 4F). We find that in both T4 and T5 the nearest-neighbour root node is most likely to be of the “Opposite” classification in a manner which is well above chance. This pattern also extends to subtype-specific pairings (Fig 4D and 4G). Here, for both T4 and T5, we see that the most probable nearest neighbour is of the opposite subtype (a to b, c to d), and the same subtype the least likely (Fig 4D and 4G; diagonal), with the orthogonally oriented subtypes positioned between the opposite and same subtypes.

This indicates a clear spatial organisation in the dendrite root node positions, highlighting the potential importance of the initial pattern of innervation before dendritic arborisation. How this result relates to the spatial position of dendrite root nodes within optic lobe columns remains to be determined however.

### Dendrite Orientation and Geometry

The asymmetric orientation of T4 and T5 dendrites has been well established [1, 2, 4, 10, 20]. Beyond this orientation however, the high resolution of these data allows for the analysis of the orientation of individual dendritic sections, and the geometry of bifurcations. We have already observed that all dendrites, regardless of subtype, occupy an elliptical space oriented along the DV axis. This alone does not show their characteristic directionality however. The manner by which individual sections within each dendrite are oriented tells us how this asymmetry may emerge, and if outgrowth during development is coordinated along the DV/AP axes, or independently.

#### Dendrite Orientation

For each dendrite we can calculate the mean angular orientation of individual sections within the dendrite, which shows a clear organisation into the four known subtypes in both T4 and T5 (Fig 5A and 5B). This dendrite-section-based analysis of orientation is in line with similar analyses conducted on the orientation of differing Strahler order sections [10]. Interestingly here the distribution of T5a dendrite orientations is, although still oriented, far less clear than all other subtypes (Fig 5B). This may relate to the previously noted more centralised dendrite root positions (Figs 2A and S1).

**Fig 5.**
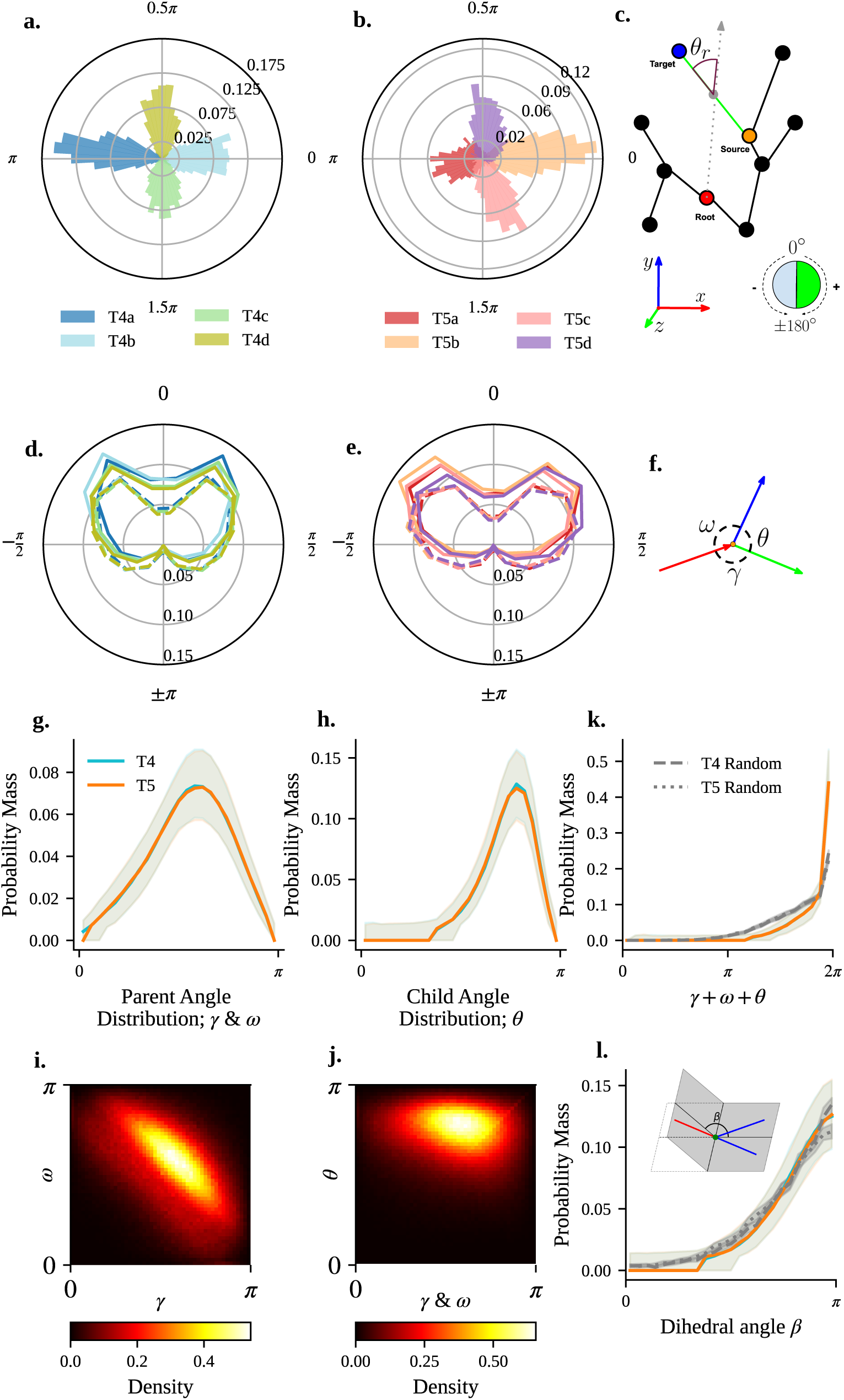
Dendrite orientation and Geometry. a) Distribution of mean section angles for each dendrites relative to the anterior-posterior axis, coloured by subtype. b) As in a, for T5 dendrites. c) Schematic illustration of radial angle calculation. Note that *θ_r_* = 0 denotes a section perfectly aligned with the ray cast from the dendrite root (gray dashed arrow). As such, segments where the source node is the dendrite root point have *θ_r_* = 0 and are excluded from analysis. Plane normal orientations used for calculation of the sign of *θ_r_* are oriented in the same direction to ensure accurate signs for *θ_r_*. d) Radar plot showing distribution of section radial angles within T4 subtypes. Solid lines show internal dendrite sections, and dashed lines denote external leaf sections. e) As in d, but for T5 dendrites. f) Schematic illustration indicating notation of bifurcation angular components. The red arrow denotes the parent section, and blue and green indicate the two child sections. *γ* and *ω* show the angles from the parent section to either of the two child sections. *θ* denotes the angle between the two child sections. g) Distribution of parent to child angles *γ* and *ω* within T4 and T5 dendrites. *γ* and *ω* are collapsed together as they are determined arbitrarily. Shaded region show asymmetric MAD within the probability mass function across individuals. h) Distribution of child angle *θ* for T4 and T5 dendrites. Line colours are the same as in g. i) Joint probability distribution illustrating inverse relationship between parent to child angles *γ* and *ω*. Data for type and subtype are collapsed together. j) as in i, showing the relationship between parent to child angles *γ* and *ω* with child angle *θ*. k) Distribution of the sum of bifurcation angular components (*γ*, *ω* and *θ*) for T4 and T5 dendrite bifurcations. Gray dashed (T4) and dotted (T5) lines show bifurcation angular component sums for randomly generated bifurcations for comparison (See Materials and Method, Geometry Analysis). l) Distributions of Dihedral angle *β*, used as a measure of 3D bifurcation flatness for T4 and T5 dendrites. Gray dotted and dashed lines facilitate comparison against randomly generated bifurcations as in k. Inset shows a schematic illustration of the calculation of Dihedral angle *β* for a single bifurcation.

We next analyse the radial orientation sections within the dendrite. In order to do so we calculate the signed radial angle of each dendrite section [21] (Fig 5C; Materials and Methods Geometry Analysis). Given the dominant elongation along the DV axis observed, coupled with four orientations of subtypes, we wish to know if either, dendrite sections align with the DV and AP axis, or alternatively, if sections orient radially outwards from the dendrite root point.

We find a tightly symmetrical distribution of radial section angles with no difference between subtypes (Fig 5D (T4) and 5E (T5)) oriented around 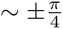. We have classified individual sections as “internal” or “external” within this analysis (See Materials and Methods, Neurons as Tree Graphs). This denotes whether a section is contained within the dendrite, starting at the dendrite root point or at a branch point, and ending at a branch point (internal), or if a section terminates at its end point (external).

We note that T5 dendrites show a broader set of radial orientations than T4 (Fig 5E in comparison to 5D), likely caused by growth in differing neuropils and different local spatial constraints. We observe that radial orientation of internal and external sections differs, with internal sections (Fig 5D and 5E; solid lines) showing tighter radial angles closer to 0*^◦^*, more closely aligned with the vector from the dendrite root in comparison to external sections (Fig 5D and 5E; dashed lines).

Although previous work has identified a radial angle distribution centred around 0*^◦^* [21], we find that within T4 and T5 dendrites, sections are still oriented consistently outward from the start of branching, however distributed symmetrically around 0*^◦^* at 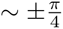. This shows that individual dendrite sections, rather than being oriented along or across the DV or AP axis, are oriented outwards from the initial branching point in a similar manner for both T4 and T5 dendrites as well as their subtypes. We also note that, although still oriented outwards, external sections show a broader distribution and are less constrained by the radial orientation away from the definite root node in comparison to internal sections (Fig 5D and 5E; solid lines compared to dashed). This indicates that the manner in which these dendrites orient individual sections, and by extension their asymmetric outgrowth, is to project outwards from the start of dendrite branching in a systematic and conserved manner across subtypes, as opposed to orienting along the DV or AP axis.

#### Dendrite Bifurcation Geometry

As our next step in building our understanding of the branching structure of T4 and T5 dendrites, we analyse the geometric structure of bifurcations in the dendrite arbourisation. Note that we limit our analysis strictly to bifurcations, although some trifurcations, or higher, are present within the dataset (∼ 6 trifurications per dendrite). Additionally, rather than calculate angles based on the whole section between dendrite root point, branch, and leaf nodes, we limit analysis to 0.1µm region around the branch node, using the full skeleton reconstruction as opposed to the reduced tree-graph representations (See Materials and Methods, Neurons as Tree Graphs; Geometry Analysis). Each bifurcation is split into three angular components γ and ω, the angles between the parent branch and each of its two children, and θ the angle between the two child branches (Fig 5F). As γ and ω are ordered arbitrarily they are collapsed together for analysis unless otherwise noted.

When first comparing the distributions of these angular components, we find no meaningfully large effects of neuron type, subtype, hemispheres, or their interactions (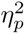 ≤ 0.01, Fig 5G and 5H). We next compare the pairwise relationships between angular components. Here we find that γ is a predictor of ω with a small effect size (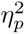 = 0.024, CI : [0.023, 0.024]), and a negative slope (β = −0.520, Fig 5I). No other effects or interactions here have an effect size greater than our minimum cut off (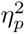 < 0.01). No such relationship is found when comparing either γ or ω with the child angle θ (Fig 5J; all 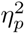 ≤ 0.01).

We next investigate the planarity of dendritic bifurcations, a property which has been observed in a number of vertebrate neurons [22–24]. Additionally, as the occupied space of T4 and T5 dendrites is naturally flat, we create a simple null model by drawing sets of three unit vectors with a parent vector oriented along the negative x-axis and the two child vectors oriented along the positive x-axis. We additionally scale the distribution of points to align with the average dispersion along each axis observed in our data for T4 and T5 separately, derived from the PC weighting (see Materials and Methods, Geometry Analysis).

First, we look at the sum of angular components for each bifurcation. The sum of angles at a bifurcation in 3D is restricted in three ways: (i) each angle will be smaller than or equal to 180*^◦^*, (ii) the largest angle will be ≤ the sum of the other two, and (iii), the sum will be ≤ 360*^◦^* [22]. The sum of angular components within bifurcations peaks at 2π, with no meaningfully large differences between types, subtypes or interaction (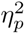 ≤ 0.01). The observed distribution is more heavily dominated by values close to 2π than observed in our random null model (Fig 5K).

Second, we compute the Dihedral angle β for each bifurcation, recommended by [22] as a measure of bifurcation flatness (Fig 5L insert; Materials and Methods, Geometry Analysis). Again, no meaningfully large difference is found between type, subtype, or interaction (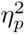 ≤ 0.01). We note that the flatness of these bifurcations is not clearly distinct from our random null model, indicating that this observed flatness is possibly a result of the relative planarity of the occupied space by each dendrites, rather than a specific result of the bifurcation process itself as suggested in [25].

Taken together, these results indicate that bifurcations within T4 and T5 are identical across both types and subtypes, with individual bifurcations being planar in nature and occupying a circular space. The inverse relationship observed between γ and ω is indicative of an optimal spatial embedding whereby both child sections are oriented forward from the parent section and maximally distant from each other. Due to the summation of angles to 2π and bifurcation planarity, the child angle θ then occupies the remainder of the circular space.

### Dendrite Topology

Our final section of analysis considers the topological and graph-theoretical structure of T4 and T5 dendrites. Analysis at this level, which is concerned with the underlying structure of dendrites, is less influenced by external factors, such as mechanistic forces dictated by their external environment. As such, we can infer from these metrics details of the internal developmental process for each dendrite, independent of external signals. Conservation of structure at this level is seen as further evidence in favour of a shared developmental mechanism across types and subtypes, independent of variability introduced between neuropils.

#### Dendrite Graph Structure

We first determine the nature of the underlying tree-graph for T4 and T5 dendrites. At one extreme, a star graph contains a single internal node and all other nodes are external, leaf nodes. For such a graph the number of branching nodes (I) is equal to 1. At the other extreme, a full binary tree is a tree-graph whereby all internal nodes have an out-degree of two. Within the reduced tree-graphs used here where all nodes with a single parent edge and a single child edge are removed, a full binary tree will have 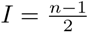 for n total nodes (see Materials and Methods, Neurons as Tree Graphs). When plotting the relationship between total number of dendrite nodes and total number of branching nodes (Fig 6A) we find that all T4 and T5 dendrites are slightly beneath this upper bound for a full binary tree. Statistical analysis identifies the total number of nodes as a predictor of total number of branch nodes with a large effect size (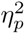 = 0.943, CI : [0.933, 0.952]) with a slope of β = 0.477. We find no meaningfully large differences between other main effects or interactions (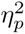 < 0.01).

**Fig 6.**
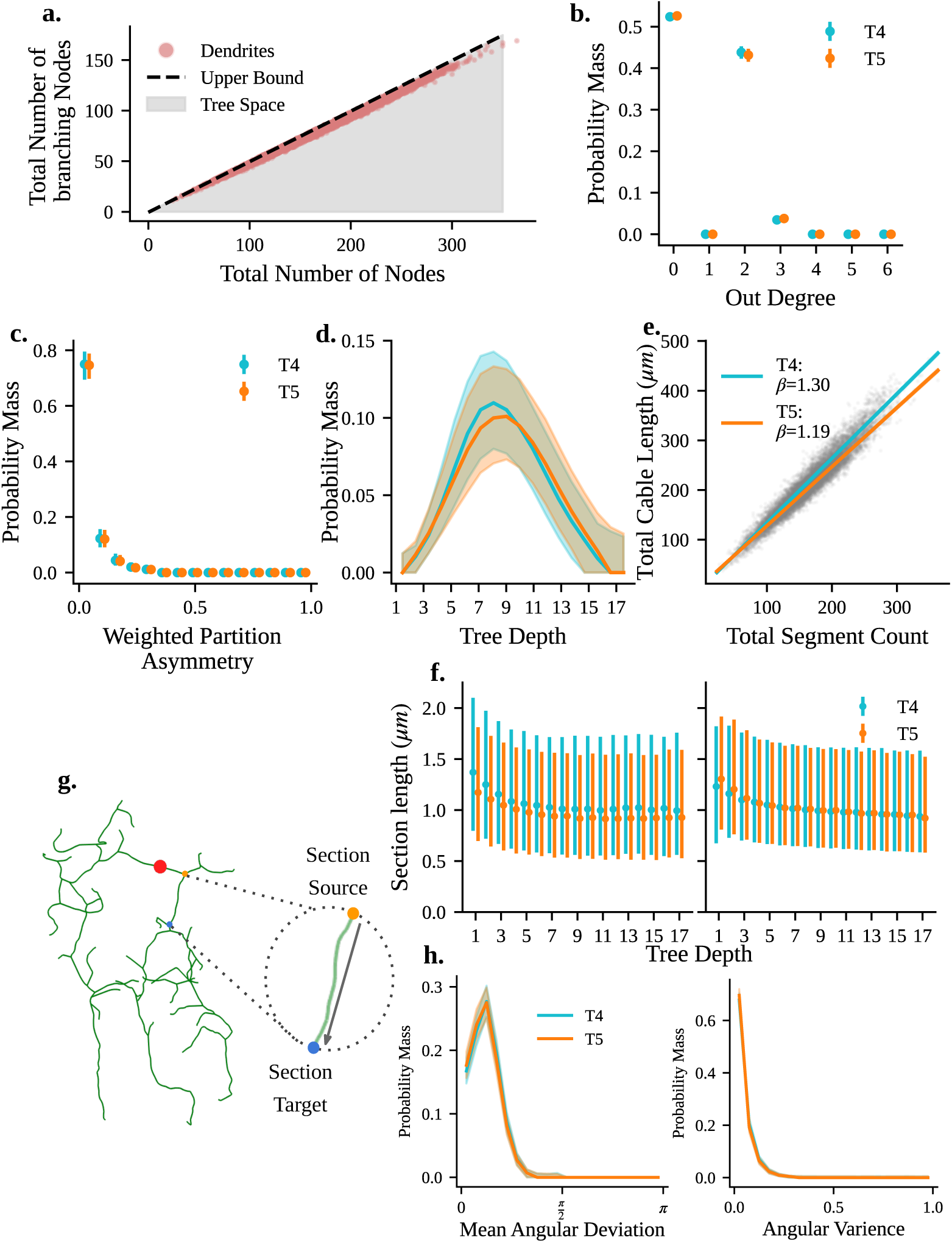
Graphical Structure of Dendrites. a) Scatter plot showing the relationship between total number of nodes and number of branching (internal) nodes within a dendrite tree graph. Gray filled in space denotes the region of possible tree-graphs, where by outside this region a given graph will no longer be a tree-graph. Black dashed line denotes a fully binary tree. b) Distribution of node out-degree for T4 and T5 dendrites. Nodes with out-degree of 0 are leaf nodes, and and out-degree of 2 show bifurcations. No nodes with an out-degree of 1 exist in the reduced tree-graph representations used (See Materials and Methods, Neurons as Tree Graphs). Error bars show asymmetric MAD of probability mass functions across individual dendrites. c) Distributions of weighted partition asymmetry across T4 and T5 dendrites (See Materials and Methods, Branching Structure and Symmetry Quantification). Values of 0 show symmetric branching and 1 show asymmetry. Error bars show asymmetric MAD of probability mass functions across individual dendrites. d) Distribution of external (leaf node) counts at increasing dendrite tree-graph depths. Error bars show asymmetric MAD of probability mass functions across individual dendrites. e) Relationship between number of dendrite sections and total dendrite cable length in T4 and T5 dendrites. f) Section length with increasing tree-graph depth for both T4 and T5 dendrites in internal (left) and external (right) sections. Error bars show asymmetric MAD of probability mass functions across individual dendrites. g) Schematic illustration showing curvature of individual segments around the euclidean vector between source and target nodes of a section. “Raw” dendrite skeletons are sampled at 0.1µm, so a single dendrite tree-graph section is comprised of multiple edges, and the euclidean distance between the source and target of dendrite sections is less than the sum of edge weights unless the section comprises of a single edge, in which case they will be equal. h) (*left*) Distribution of the mean angle of edges within a section relative to the euclidean vector between source and target within the same section. Error bars show asymmetric MAD of probability mass functions across individual dendrites. (*right*) Angular variance of edges within a section relative to the euclidean vector between the source and target point of the same section. Error bars show asymmetric MAD of probability mass functions across individual dendrites.

When observing the out-degree distribution of dendrite tree-graphs we see that these dendrites are dominated by nodes with an out-degree of 0 (all leaf nodes) and 2 (bifurcating branch nodes). Interestingly, there is a highly consistent but small number of trifurcations within the dataset (∼ 6 per dendrite). We note that these trifurcations could be a result of the skeletonization process, although their consistency leads us to believe otherwise. We find no meaningfully large differences between neuron types, subtypes, hemispheres, or interactions (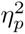 ≤ 0.01).

We next consider the symmetry of dendrite tree-graphs. Tree asymmetry has long been used as a metric in the analysis of neuron morphology [26–28]. Symmetry here refers to the symmetry of branching through the dendritic tree and is calculated at each branch node. As an example, if one pictures a bifurcation whereby the left downstream node is a termination, and the right contains the entirety of the downstream subtree which follows this same right-dominated rule, this bifurcation would be asymmetric and as such have a tree asymmetry value of 1. On the other hand, if all branching nodes downstream from a bifurcation are also bifurcations, this subtree would be symmetric, and as such have a tree asymmetry value of 0. We implement here a weighted tree asymmetry metric, similar to [28] (See Materials and Methods, Branching Structure and Symmetry Quantification). Rather than using a single summary statistic per dendrite, we show the full distribution of scores across a single dendrite, showing the clear symmetric nature of T4 and T5 dendrites (values close to 0 are closer to symmetric, Fig 6C). This is in line with previous observations [26, 28, 29] of tree symmetry in neuron morphologies. Again, no meaningfully large differences are found for neuron type, subtype, hemispheres or interactions (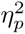 ≤ 0.01).

We note that external leaf nodes are distributed throughout tree graph depths (Fig 6D), again with no differences observed between type and subtypes (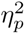 ≤ 0.01). Much like the relationship between total dendrite cable length and dendrite convex hull volume, and between dendrite convex hull volume and the number of sections within a dendrite (S2 Fig C and F), there is a strong relationship between the total dendritic cable length and the number of individual segments within each dendrite (Fig 6E). With the number of segments as a covariate of total cable length we find segment count is a strong predictor of total cable length (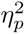 = 0.509, CI : [0.463, 0.561]). With a slope of β = 1.357 this tell shows that all dendrite sections are, on average, 1.357µm in length.

Together, these results speak of a highly conserved topological structure for all T4 and T5 dendrites, as well as their subtypes. We do not find any distinguishable characteristic within this structure which would be in favour of arguing for differences between subtypes mediated by differing orientations, or neuropil specific differences between T4 and T5.

#### Dendrite Section Lengths

We next turn to investigate any differences in the individual sections between T4 and T5 and their subtypes. As was done previously when considering the radial orientation of individual sections, we again include the internal or external nature of individual sections within our analysis. We first investigate the decay of section lengths as a function of tree depth (Fig 6F). Tree depth here refers to the number of (unweighted) steps taken from the root of the dendritic tree. We find no meaningfully large differences between hemispheres, neuron type, subtype, source, or interactions (all 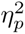 ≤ 0.01). We note, however, that sections closer to the dendrite root (with lower depth), particularly within internal sections, appear marginally longer before the section lengths stabilise. This observation of an initial decay followed by stabilization has been observed previously in mouse Purkinje cells [30], and in line with an optimal wiring and conduction time trade-off, which would imply proximal longer runs and distal finer segmentation [31]. T4 internal section lengths also appear qualitatively greater than T5 across all tree depths (Fig 4F; *left*).

The individual section within a dendrite are a collection of short edges which curve around the individual dendrite root point, branch and leaf nodes within the tree graph (Fig 6G). We would like to understand how much deviation there is from the shortest vector between points as the dendrite section projects between them. As such, we measure for each section the mean angular deviation between individual tree graph edges and the euclidean vector between the source and target point of the section, and the angular variance of the individual edges. We find no differences between types and subtypes (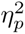 ≤ 0.01), showing the curvature within individual sections is tightly conserved again across hemispheres, neuron types, subtypes, and sources. Additionally, individual sections show very little deviation, indicating that outgrowth is essentially straight for within all dendrites between branching and terminal events.

We next investigate the distribution of individual section lengths across T4 and T5 and their subtypes. As well as “raw” section lengths, we additionally scale individual dendrites along their internal axis to have equal variance, creating uniform spherical dendrites without changing the topological structure. The intention of this scaling is to remove the influence of the local dendrite environment, such as external mechanistic forces, or the confines of the neuropil layer from the length of dendrite sections, without altering the topological structure. Ideally, this results in a dendrite representation independent of external influence (Fig 7A and 7B).

**Fig 7.**
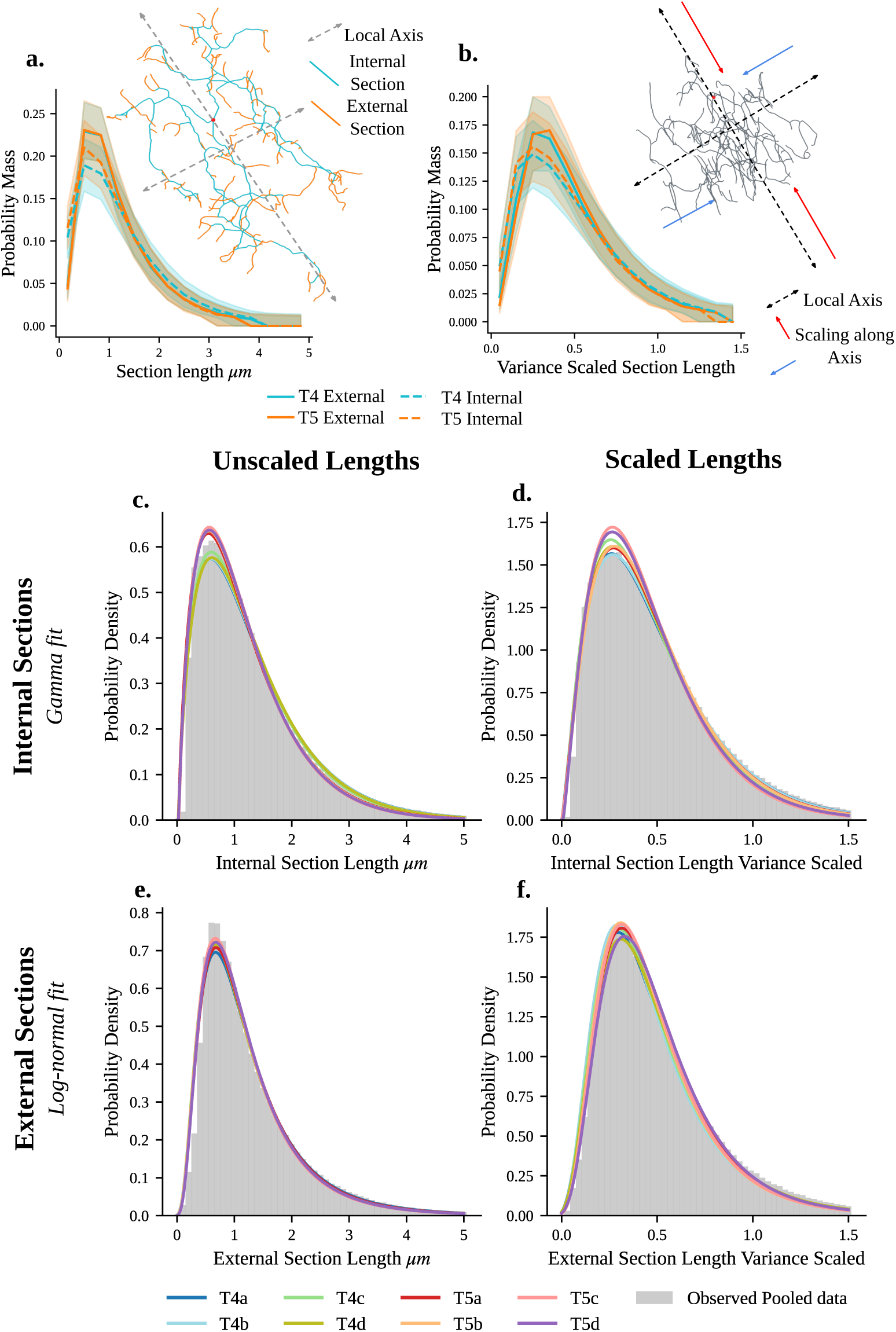
Dendrite Section length Distribution and Explorative Distribution Fitting. a) Distribution of dendrite section lengths for T4 and T5 dendrites, with dashed lines showing internal sections and solid lines showing external leaf sections. Schematic (insert) shows example neuron with internal (cyan) and external (orange) sections coloured for illustration, along with first two internal axis. Errors show asymmetric MAD of probability mass functions across individual dendrites. b) Distribution of section length as in a, but for variance scaled dendrites (See Materials and Methods, Alignment and Scaling). Dendrites are individually scaled to have equal variance along each internal axis in order to account for neuropil specific differences. Schematic (insert) illustrated variance scaling along internal axis. This results in a spherical dendrite. Errors show asymmetric MAD of probability mass functions across individual dendrites. c - f) Best fitting probability distribution for pooled section lengths within each subtype. Left hand plots show unscaled section lengths (c and e) and right hand plots show variance scaled section lengths (d and f). The top row shows internal sections (c and d) and bottom external sections (e and f). Both unscaled and variance scaled internal section lengths are best fit by a gamma distribution (c and d). Both unscaled and variance scaled external section lengths are best fit by a log-normal distribution (e and f). Gray histograms show the probability density for all pooled data.

Interestingly, in both original unscaled and scaled datasets, we find no meaningful differences between hemispheres, neuron types, subtypes, source, or their interactions (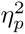 ≤ 0.01). Due to the high skew observed in these data, we also test log-transformed values, which have the same result (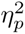 ≤ 0.01). Our analysis here again shows a highly conserved structure with no meaningfully large differences between T4 and T5 dendrites and their subtypes.

#### Explorative Analysis of Section Length Distributions

Previously, measures of dendrite and axon section lengths have been observed generally with a range of right-skewed distributions [28, 32–36]. However, understanding the specific nature of the observed right-skewed distribution tells us something of the underlying mechanistic process. For example, models of multiplicative dynamics have been shown to directly give rise to the observed log-normal distribution of spine size in the auditory cortex of mice [36].

In order to better understand the specific distributions observed here, we perform an exploratory analysis to determine which from a set of six distributions best fits the observed section length data. We choose to compare exponential, log-normal, gamma, Wald, log-logistic, and minimum Weibull distributions. Although we note that an exponential distribution is a special case of the gamma distribution with k = 1, we keep the two distributions distinct for clarity. Each of these distributions is an example of a right-skewed distribution, and can be related to differing generative processes. We fit all six distributions to each type, subtype, and source (internal vs external section) combination for both raw and variance scaled section length data. Distributions are compared using the Bayesian Information Criterion (BIC). Interestingly, there is always a single consistent best fitting distribution for types and subtypes. Additionally, the best fitting distribution is consistent within both scaled and unscaled dendrites.

For internal sections, data is consistently best fit by a gamma distribution (Fig 7C and 7D). For raw section lengths, fitted distributions show a slight distinction between T4 and T5 (Fig 7C), which is also in line with the small qualitative difference observed in internal section lengths as a function of tree depth (Fig 6F). This difference is no longer present in variance scaled section lengths, however each group here shows greater variability in comparison to other fits, although there is no clear structure to this variability (Fig 7D). For external section lengths, we find data is consistently best fit by a log-normal distribution for all types and subtypes, with little variability between fitted distributions (Fig 7E and 7F).

Although the first place rankings are consistent within all groups examined, this explorative analysis does not represent a definitive analysis. Although n is exceptionally large within this dataset (∼ 250000 segments per fit), distinguishing statistically between these similar distributions is a challenge [37, 38]. When considering the range of BIC scores for each fit, we note that their is significant overlap between subtype groups (S5 Fig), however rankings are still consistent. As such, we would encourage caution in over-interpreting these results. However, we would argue that this result is evidence in favour of fundamentally differing generative processes underlying branching rates and termination rates.

## Discussion

By leveraging the high resolution morphological data available through new EM reconstructions of all T4 and T5 dendrites within a single *Drosophila* brain, we are able to offer a quantitative characterisation of the similarities and differences between T4 and T5 dendrites, as well as their subtypes, at a level not available through typical imaging techniques. In doing so, we have aimed to shed light on specifying the features relating to these dendrites characteristic directed spatial embedding, in order to understand the origins of this symmetry breaking.

### Dendrites are Structurally Homogeneous

Given the distinct characteristic orientation of T4 and T5 dendrite subtypes [3], one may assume that this would lead to morphological differences within the dendrite branching structure. We, however, observe consistent homogeneity in all geometric, and graph-theoretical, measures across all T4 and T5 dendrites, as well as their subtypes, across both hemispheres.

The dendrites of T4 and T5 neurons, and their subtypes, are all well characterised by the same close to binary and symmetric tree-graph with little variability across hemispheres, neuron types, and subtypes (Fig 6A to 6C). Interestingly, the deviation from a binary tree manifests in a small but consistent proportion of trifurcations (Fig 6B). Their consistency leads us to believe that they are less likely to be a direct result of the skeletonization procedure. This may indicate that they are biologically meaningful, for example, being Steiner points which minimize the total dendritic cable [39], however further analysis is needed in order to determine if this is the case.

We also observe a striking similarity in the geometry of dendritic branching, even though individual subtypes are globally oriented (Fig 5). The angular components of all dendrite bifurcations are indistinguishable, as is their planarity, illustrating that dendritic branching itself is indistinguishable between hemispheres, neuron type, and subtype. The implication here being that the mechanism of branching within these dendrites is conserved and, regardless of the ultimate directionality of the individual dendrite, or the neuropil which it exists in, the result of a consistent mechanism.

The observed planarity is potentially the result of additional constraints on branching geometry. Previous work has related the flatness of bifurcations with both the stochastic process of connecting to the closest point in space randomly [25], and being an emergent property of the local 3D elastic tensions in neurites, where distal parts of branches develop force equilibrium when anchored to the substrate [22]. Either mechanism is plausible here, although we cannot disregard that the observed flatness is a result of the relative planarity of the occupied space of these dendrites.

Further, our results indicate a simple rule whereby the two child sections are maximally separated from each other. This follows from the relative planarity of these bifurcations, coupled with the negative correlation between both parent and child branching angles. This maximal partitioning may relate to self-avoidance mechanisms, as well as optimal spatial embedding.

These dendrites also show similar spatial occupancy, shown in their elliptical structure (Figs 2A and S1), their consistent elongation along the Dorsal-Ventral axis regardless of subtype (Fig 2C), as well as neuropil layer depth innervation (Fig 3). Once appropriate scaling is applied, the mean volume of these dendrites additionally does not differ (Fig 2G), in line with the importance of the spatial occupancy of ∼ 7 columns and necessities of motion detection to integrate across multiple adjacent visual points [2, 4]. Independent of this spatial scaling, the total cable length and number of sections within the dendrite tree-graph also does not differ markedly between hemispheres, neuron types, or subtypes (S2 Fig, S1 Text: Statistical Analysis of Dendrite Size Metrics).

Together, these analysis highlight that the spatial occupancy, geometry and topology of these dendrites is, essentially, identical. From this, it seems unlikely that differing mechanisms of branching, or space filling, are employed during the emergence of these dendrites asymmetry.

### Observed Differences are Exclusively Constrained to Spatial Embedding, Not Structure

Although the structure of individual dendrites is highly conserved across types and subtypes, there remain some insightful differences particularly focused on elements of their individual spatial embedding. Of particular interest is the differing distribution shape and spatial organisation in all size based metrics. Specifically, the population of T4 dendrites shows a heavy tail not present in the comparably symmetric T5 distribution (Figs 2F, S2A and S2D). The tail of larger T4 dendrites is focused within the anterior-ventral region of Medulla layer 10 (Figs 2H, S2B and S2E). There are two opposing explanations to this observation. Either, the visual columns within Medulla layer 10 are physically larger, or column size is uniform and these larger dendrites, by extension, sample from a greater proportion of the visual space. Interestingly, it has been previously observed that within the compound eye, across *Drosophilidae* species, larger lenses are present within the anterior - ventral region [40–42]. Strikingly, we find that the column space within Medulla layer 10, which determines a T4 dendrites sampling of visual space, is uniform (S3 Fig). This indicates that individual T4 dendrites within the anterior-ventral region span across a larger portion of the columnar grid, and as such seem to integrate visual inputs from a larger portion of visual space. It is important to note that this this only relates to the spatial span across the columnar space in relation to Mi1 morphology, and does directly not relate to difference in features such as input density, or the spatial distribution of synapses. That this region of enlarged dendrites correlates with larger lens sizes on the retina seems unlikely to be a coincidence. Importantly, future functional analyses will be needed to determine if there is any implication to this observation in synaptic integration of receptive field size, or if this is merely a quirk of morphology.

Interestingly, we also find evidence of differences in the positions of the initial point of branching within these dendrites. Within both T4 and T5 neurons, there is some evidence in favour of subtype specific variability in the depth of dendritic root points within their neuropil layer. Statistically, this is dominated by the ‘a’ subtypes being closer to the layer surface. Qualitatively, particularly within T4 dendrites, our analysis indicates a possible split between horizontal and vertical subtypes when considering layer depth for dendritic root points (Fig 3C). Interestingly, within the left hemisphere, T4 dendrite root point depths are significantly more varied in comparison to the T4 dendrites within the right hemisphere, or T5 dendrites in general. As individual dendrites are reviewed in a random order across the mix of all subtypes and hemispheres, this is unlikely to be a systematic bias introduced within our pre-processing procedure. We cannot rule out that this is some artefact within the reconstructions of T4 dendrites within the left hemisphere, but these have also been independently proofread within the FAFB-FlyWire dataset.

We also observe differences between horizontal and vertical subtypes in terms of the dendritic root point. We see a clear difference between nearest-neighbour partners in T4 horizontal and vertical subtypes when considering oppositely oriented nearest-neighbours, whereby horizontal subtypes are closer nearest-neighbours than vertical (Fig 4B). This trend is not present in T5 neurons (Fig 4E), and is not a result of the observed differences in the depth of dendrite root points (S4 Fig). This observed difference may be a result of the columnar structure itself, as noted in [4], whereby the columns within both layers are elongated along the DV axis, however within Lobula layer 1 they are additionally shortened along the AP axis, resulting in a triangular columnar array as opposed to hexagonal. When we consider the aspect ratio (AR) between AP and DV axis between neuropils (Simply AP / DV extent within each neuropil) we find the Lobula is, indeed, proportionally narrower (AR: Medulla right = 0.66, Medulla left = 0.72, Lobula right = 0.55, Lobula left = 0.58). How dendritic root point innervations into their target neuropils map to the columnar lattice is unclear, for example it is unclear if the dendrite root point of each subtype within a column are all centrally located, or some sit nearer the boundary. The narrower columns along the AP axis within the Lobula may serve to constrain horizontal and vertical subtypes, forcing them closer together in comparison to horizontal and vertical subtypes within Medulla layer 10. Additionally, as the first NB division during development separates horizontal and vertical subtypes, the manner of targeting used to innervate their eventual neuropil prior to dendritic outgrowth may differ, facilitating the observed difference. As a final consideration, Medulla layer 10 and Lobula layer 1 also differ in some manner in terms of their global structure. Medulla layer 10 is somewhat closer to a slice taken from the surface of a sphere, whereas Lobula layer 1 shows greater variability and deviations from this, although still well approximated by a section taken from a spherical surface. These differing geometries neuropil can also have an impact on our observations of differing elements of spatial embedding within these dendrites.

Of additional interest is the observed subtype, but not type, specific relationship in the extent of dendrite innervation along the dorsal-ventral (DV) and anterior-posterior (AP) axes. Vertical (c,d) subtypes show a greater elongation along the DV axis, and narrower width along the AP axis, a trend which is smaller within the horizontal (a,b) subtypes (Fig 2D). Interestingly, however, the difference between elongation along the DV and AP axis is negatively correlated (Fig 2E), indicating that as each dendrite extends along the DV axis, it extends less along the AP axis proportionally. This could be evidence of a cost-constrained mechanism limiting the total size of dendrites, coupled with dominant, subtype-specific, elongation along the DV axis. The general elongation along the DV axis is in line with observed column elongation along this same axis [4], however how or why the extent of this elongation is subtype specific is unclear.

These results highlight subtle differences between the T4 and T5 dendrites. Notably, in terms of the initial innervation pattern, we see evidence of a horizontal / vertical subtype split within T4 and the Medulla which is not evident within T5 and the Lobula. It is interesting that we only see this separation between horizontal and vertical subtypes in relation to the point at which the dendrite starts its branching. Given that these dendrites first innervate their entire target neuropil, whilst remaining unicolumnar, before asymmetry emerges within the dendrites, how this may play a role in the emergence of dendritic orientation, and what impact the horizontal / vertical distinction may have in T4 but not T5 is unclear. What is clear, however, is that regardless of the starting position of dendritic branching, the resulting structure of the differing dendrites for both T4 and T5, and their subtypes, is conserved.

Second, the observation of larger T4 dendrites alongside a uniform columnar space, does raise the interesting question of how this may impacts function. We have shown that within the region where lenses on the retina are larger, T4 dendrites additionally appear to span a larger area, even though the columnar grid through which they sample their visual inputs, is uniform. This directly implies that T4 dendrites within the anterior-ventral region integrate motion over a marginally larger region of visual space, and as such would be expected to posses a larger receptive field. We are not aware, however, of any functional studies which show this. On the other hand, given the fundamental necessity of motion detection within visual processing, this may well be an artefact of these dendrites development which is modulated by functional tuning, leading to a morphological, but not functional, difference.

### The Emergence of Orientation

Given the homogeneity of dendrite structure across subtypes and types, it is important to pinpoint exactly what is meant by their characteristic orientation. At some stage between 36 and 72 hours APF [8], the well documented orientation of these dendrites must emerge [1, 4, 10]. We have shown that these dendrites, regardless of their spatial orientation, share a common structure. Their spatial embedding, although more variable, follows the principle of elliptical space filling elongated along the DV axis. Additionally, we have shown that, although the directed subtypes can be distinguished by the mean orientation of section angles relative to the start of dendritic branching, these sections are consistently oriented away from this point. This appears indicative of a shared space filling algorithm across these dendrites regardless of subtype specific orientation [43, 44].

The characteristic orientation of these dendrites then seems to be determined by the centrally offset root position of the dendritic branching process in relation to the centre of the occupied elliptical space (Figs 2A and S1). How then, given that individual sections are oriented radially away from the dendrite root point (Fig 5D and 5E), does the mean direction of these sections result in orientation? To illustrate a process by which orientation emerges as a result of offsets in the position of initial branching (the dendrite root point) relative to the final filled elliptical space, consider an ellipse in which a point is randomly placed at an off-centre location. If short lines a drawn within the ellipse, oriented radially away from the point but directed towards the boundary, the average angle of these lines relative to a global reference will be oriented, as is the case between subtypes here. It is evident that the initial branching positions for all subtypes exhibit a consistent offset relative to the occupied space. Specifically, dendrite roots in the vertical subtype are positioned centrally within the lower and upper regions of their spanned area, whereas horizontal subtype roots are located within the central left or right regions (Figs 2A and S1). This configuration enables dendritic orientation within the occupied space, while preserving the structural homogeneity independent of overall orientation.

T5a dendrites, however, show a markedly more central dendrite root point (Figs 2A and S1). Interestingly, the orientation of dendrites from the mean angle of dendrite sections for T5a is still present, however it possesses a somewhat broader and less directed distribution in comparison to all other subtypes (Fig 5B). While this observation is in line with the above described mechanism: a more central initial position would result in reduced directionality, it is unclear why this is the case within T5a dendrites. Similarly, within T5c dendrites we see a shift in their orientation toward the posterior direction (Fig 5B), in line with the observed shift of the dendrite root point towards the anterior (Figs 2A and S1). This is not observed however within the T5d subtype, although we would expect it to be. Although we observe this result separately within both hemispheres, we are cautious to not rule out the possibility of artefacts within either the FAFB-FlyWire dataset itself or our processing pipeline, although these seem unlikely. Future work should seek to validate, or refute, this observation in a separate dataset before speculating about the broader impact of this observation.

### Insights into Dendrite Development

We have set out our analysis in order to quantify the morphology of all dendrites within these closely related populations of neurons with the aim of better understanding how the characteristic subtype specific orientation may develop between 36 and 72 hours APF [8]. Our analysis has shown that these dendrites are structurally homogeneous, which we take as strong evidence of a shared algorithm of dendritic outgrowth within this period. Our analysis indicates a developmental process whereby new dendrite arbours orient outwards from the initial branching point to occupy space in an optimal manner at equivalent rates (Fig 6) and with shared geometry within both T4 and T5, and their subtypes (Fig 5). This, alone, would produce a circular dendrite, and it is unclear how such outgrowth facilitates symmetry breaking within this period.

A likely contender for the emergence of orientation during development is that subtypes make use of chemical gradients. The identification of chemical gradients within the Medulla and Lobula during development which may be related to T4 and T5 development have remained elusive however. One contender would be the identified DWnt4 and DWnt10 A/P gradient identified within the Medulla during development [45]. However this gradient is observed at 25 hours APF, and not present at 45hrs APF. This is an issue, as the dendritic orientation of T4 and T5 dendrite subtypes does not start to emerge until later than 36hrs APF [8].

Given the absence of a definitive chemical gradient, we propose two potential local cues which could influence dendritic orientation in light of our analysis which would maintain structural conservation. Our analysis shows that dendrites innervate their target neuropil in a subtype specific manner, with opposingly oriented dendrites positioned as nearest neighbours (Figs 4 and S4). Previous work suggests that these nearest neighbour partners likely originate from the same neuroblast (NB) [7]. Additionally neurons from the same NB have been shown to be able to distinguish themselves from other neurons from the same lineage [46, 47]. During the initial dendritic outgrowth phase where dendrites remain confined within a single column [8] dendrites originating from the same NB may provide attraction or avoidance cues determining later orientation and outgrowth. Second, each dendrite densely overlaps in space. During development, neurons will form synapses promiscuously and, taken out of their developmental context, form any synapses they can [48]. This implies that dendrites will compete spatially for synapses. For two dendrites which initiate branching close together, this competition will be reduced in the opposing direction, facilitating directed outgrowth.

While such promiscuity could disrupt the precise synaptic compartmentalisation observed within these dendrites [4], we have shown that they orient across, not along, the neuropil depth (Fig 3D and 3E). If input partners are stratified within the neuropil layer, or if their innervation times differ, spatial compartmentalisation of synapses may, in part, be a result of the spatial or temporal availability of these partners during development. This allows for promiscuity during development, while maintaining the spatial compartmentalisation of synapses onto these dendrites.

Although we invite caution in putting to much emphasis on the specific distributions identified within our explorative analysis of section lengths, the observed distributions do offer some interesting interpretations to development. We observe a gamma distribution as the consistently best fitting distribution to internal dendrite section lengths, effectively the inter branch interval. The gamma distribution relates to a process involving the necessity for multiple events to occur before a branching event takes place. Giving a more concrete example, this implies that a branching event will occur during growth after *n* sub-events, which are exponentially distributed, have first occurred [49]. When branching is treated as a Poisson process, and multiple events are required before a stable branch occurs, we would expect to observe a gamma distribution as a result [50]. Empirically fit models which treat branching in this manner have previously been used to accurately capture the observed distribution in real data [51–53]. This assumes a stable dendrite branch requires some number of molecular sub-events before formation. Actin nucleation, specifically transient, localized Arp2/3 recruitment, precedes branch formation in *Drosophila* DA neurons [54], as an example of one such potential molecular sub-event. Further, microtubules have been shown to transiently invade branch precursors, providing a pathway for additional material required for stabilisation, initiating changes in dendrite spine morphology [55]. MAP2c in cultured hippocampal neurons has also been shown to promote stabilisation of neurites, through the coupling of F-actin and microtubules, consolidating stabilisation [56]. Our data is indicative of a complex multi-event mechanism which governs branching within these dendrites.

Within external dendrite sections, a log-normal distribution is the consistently best fitting distribution to the data. A log-normal distribution has been previously related to dendrite spine outgrowth and lengths [36], as well as axonal tree section lengths [28]. The log-normal distribution is a result of a central limit theorem-like process resulting from the product, rather than the sum, of multiple samples [57, 58]. Dendritic spine lengths have previously been observed to follow this type of distribution, both experimentally and within modelling studies [36, 59, 60]. How this may relate mechanistically to the outgrowth of dendrite external sections is not as clear however. Live imaging has shown stochastic switching between growth, paused, and shrinking states in an episodic manner within dendrite tips [61]. This dynamic state switching, coupled with the available resources in the dendrite tips, could, in principle, generate the multiplicative effects required to produce the observed log-normal distribution [28].

### Future Analysis and Perspectives

Multiple avenues of future work open up in order to better understand T4 and T5 dendrites from connectomics data. The most obvious is to investigate comparative differences in T4 and T5 dendrites between datasets, utilising the available male optic lobe connectome which has been recently released [62]. This would allow us to better capture variability across individuals, and also allow for sex-specific comparisons relating to differing size, shape, and orientation of *Drosophila* compound eyes between males and females across species [41].

Second, it is important to understand the synaptic distributions within T4 and T5. Synaptic inputs are vital to dendritic development [63–65], and given the fundamental role of motion detection within visual processing, it seems likely that the functional output of T4 and T5 dendrites is tightly regulated. Further, the origin of the distinct spatial compartmentalisation of synapses onto T4 and T5 dendrites is not well understood. Extending our analysis to included synaptic information, as well as additional functional imaging or modelling with the available highly detailed dendritic morphology, will be insightful.

Finally, we have limited out analysis of columns to only the size of columns as defined by Mi1 neurons. As we consider the optic lobe column structure to be the map of visual space through which these dendrites perform motion detection, it will be important to extend these analysis to better understand the relationship between visual columns and the spatial embedding of these dendrites, particularly in light of the observed size differences of T4 dendrites within the anterior-ventral region. However, see [10, 66] for analysis of T4 and T5 neurons which includes visual space mapping. Of additional interest will be understanding how the structural relationship between the start of dendritic branching is related to the column map, and if this relates to emergent orientation of these dendrites through the selection of columns which they ultimately sample from.

### Limitations

The data presented here are from a single adult female *Drosophila*, offering a view of a single snapshot of T4 and T5 dendrite morphology. Although thought of as complete, and offering a new gold standard in data quality, some noise is undoubtedly introduced from the complex processing steps needed to generate a usable dataset from the EM image stack.

Before consideration for inclusion within this study, we ensure that all dendrites are labelled as complete and proofread within the FAFB-FlyWire dataset, ensuring they have already been independently reviewed before our use. We additionally manually verifying the accuracy of each individual dendrite. However, particularly small branches are likely missing from the morphological reconstructions of neurons. It is worth noting however that previous work using manually reconstructed dendrites, the gold standard in morphological reconstruction, shows that such small branches are not directed and typically concentrated on connecting a small number of synapses to the dendrite [10], and a such minimally impact the dendritic morphology.

In addition to this we would argue that as this error is likely missing at random, and as the *n* of T4 and T5 neurons is sufficiently large, this error rate is likely uniform across the dataset and is unlikely to influence our core findings. As these data also come from a single *Drosophila*, it is also possible that different individuals show differing results, keeping in mind however that our results are consistent across both hemispheres. Given the fundamental nature of T4 and T5 neurons in early visual processing and motion detection, we feel this is unlikely to be the case. The question of individual variability between datasets is an intriguing one which requires further work and comparative analysis between datasets in order to be fully understood.

### Conclusions

In summary, we show that the underlying structure of T4 and T5 dendrites, as well as their subtypes, is indistinguishable. There are however, some important type- and subtype-specific differences in the manner of spatial occupancy and initial dendritic branching. With this detailed characterisation of T4 and T5 dendritic morphology, and given the observed homogeneity, we conclude that the underlying algorithm of dendritic outgrowth which leads to orientation between 36 and 72 hours APF is likely conserved across all T4 and T5, and their subtypes. However, the specific mechanisms underlying these dendrites’ characteristic orientation cannot be clearly identified here.

## Materials and Methods

### T4 and T5 Dendrite Annotation and Pre-processing

The FAFB-FlyWire dataset comprises of ∼ 140000 fully reconstructed neurons from a single 4 × 4 × 40nm resolution electron microscopy (EM) image stack [13–16, 67]. Neuron type annotations within the right optic lobe of this dataset are available through [9], and include ∼ 6000 annotated T4 and T5 neurons. These annotations have been updated since the publication of [9] to include both optic lobes bringing the total number of annotated T4 and T5 neurons to ∼ 12000. Neuron type annotations use the current annotation state within FAFB-FlyWire as of April, 2026 [9, 14–16].

Each neuron is represented as a 3D surface mesh. Skeletonization, the method of extracting tree-graph representations from neuron surface meshes, is implemented using a standard procedure [68, 69]. The correct rooting of each neuron at the soma is manually verified and corrected if needed. All reconstructions have been annotated as complete within the FAFB-FlyWire dataset. After skeletonization, all neurons are converted to µm from nm, and resampled to 0.1µm. Table A in S1 Text gives counts by subtype and hemisphere for available and included dendrites.

With skeletonization, each neuron is modelled as a tree-graph (See Materials and Methods, Neurons as Tree Graphs). We further require the accurate annotation of the dendritic compartment of each neuron tree-graph in order to extract and analyse dendrite morphology (See Materials and Methods, Dendrite Extraction and Pre-processing).

### Neurons as Tree Graphs

Each neuron represented as a tree-graph is rooted at the soma, in the case of complete neurons, or the first branching point of nodes within the dendritic subtree (referred to as the dendrite root point), and all edges are oriented away from this root node. For completeness, we provide a brief overview of graph-theory specific to directed tree-graphs here and the components relevant to this manuscript.

A neuron is modelled as a directed tree:

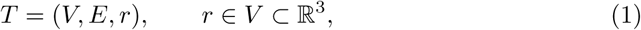

where *V* is the set of 3-D coordinates defined by the skeletonization algorithm and *E* ⊂ *V* × *V* the set of oriented edges. *r* denotes the root node of the tree. *e* = (*u*→*v*) denotes an edge from *u* to *v*, where *u* is the source node, and *v* the target.

Each edge carries the Euclidean length *w*(*e*) = ‖*v* − *u*‖_2_. The tree weight (total cable length) is:

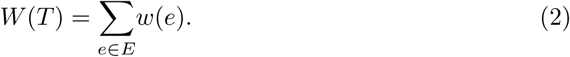

For *v* ∈ *V* :

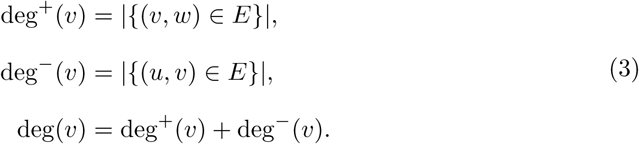

Where deg^+^ denotes the out-degree of a node, deg*^−^* denotes the in-degree, and deg(*v*) the total degree. As edges point away from the root, deg*^−^*(*r*) = 0, and deg*^−^*(*v*) = 1 for every *v* ≠ *r*. branch nodes have deg^+^ ≥ 2 and leaf nodes deg^+^ = 0. An edge is *external* if *e_v_* is a leaf, otherwise *internal*.

For any two nodes *u, v* ∈ *V* there exists a unique undirected path

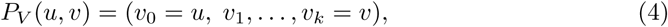

with *k* = *d*(*u, v*) steps.

The corresponding edge sequence:

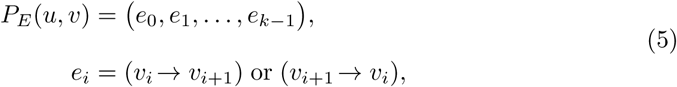

lists the *k* oriented edges traversed in order; note that orientation may be reversed relative to *E* when the walk proceeds toward the root.

Unweighted and weighted path lengths are

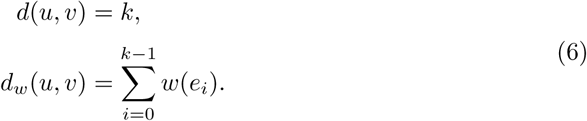

Node depth is its unweighted distance to the root, *h*(*v*) = *d_w_*(*r, v*), and an edge inherits the depth of its distal node.

Every maximal path of intermediate nodes with deg^+^(*v*) = deg*^−^*(*v*) = 1 is contracted into a single edge whose weight is the sum of the collapsed weights. The reduced graph is *T_r_* = (*V_r_, E_r_*). Such reduced trees dramatically minimise the number of nodes and edges within each neuron skeleton.

For any node *v* ∈ *V* we define the *subtree* rooted at *v* as:

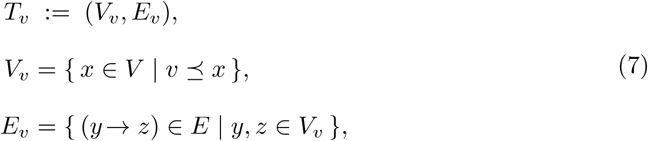

where *v* ⪯ *x* means that *v* lies on the unique root–to–*x* directed path in *T*. Thus *T_v_* contains *v* and all its descendants together with the edges that connect them, and is itself a rooted, oriented tree with root *v*.

### Dendrite Extraction

Previously, separation of neuron morphologies into biologically relevant subtrees, such as the axon and dendrite, has been done manually [70] or derived from synapse data [71, 72]. Within this dataset of ∼ 12000 neurons, manual annotation of each dendrite is not practical. Methods based on the input-to-output ratios of synapses [71, 72] assume clearly defined information flow between compartments, which does not always hold [70], and require additional synapse annotations. We instead implement a novel, semi-automated method to annotate the dendritic compartment of T4 and T5 neurons based on tree structure alone.

Given a tree-graph correctly rooted at the soma, we then identify the subtree in *T* which maximises the number of leaves within the subtree whilst minimizing the sum of edge weights using a simple metric, Φ:

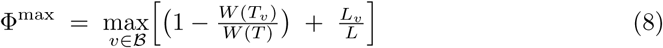

Where *B* is the set of branch nodes within *T*. Although often the identified subtree will be the dendrite within T4 and T5 neurons, due to either routing errors resulting from the surface mesh and skeletonization procedure, or instances where no single node isolates the root of the dendritic subtree, we manually verify and modify the identified dendritic compartment for all neurons, and exclude those with an inaccurate or visibly incomplete dendritic tree.

This manual verification step is completed using the GUI available within the NeuRosetta toolbox (See Materials and Methods, Data and Code Availability). This GUI allows for the re-rooting of neurons, manual subtree annotation from a single point (alongside automated identification using the above metric) and flagging of neurons for exclusions from further analysis. This manual review step can be completed using this GUI in a relatively short time by a single expert reviewer.

### 0.1 Global Population Dendrite Alignment

As T4 and T5 dendrites exist in two separate neuropils, across two hemispheres, we perform a global alignment step in order to ensure each dendrite population is aligned with the dorsal - ventral and anterior - posterior axis in visual column space. This process is performed by taking the following steps, where a dendrite population refers to each individual population of dendrites within each hemisphere, so for example all T4 dendrites within the right hemisphere.

First, we subtract the mean coordinate within each population, and perform a rotation. The aim of this rotation is to ensure that the point cloud is oriented in a way which ensures that the longest extent of dispersion is along the y-axis, the second along the x-axis, and third along the z-axis in euclidean space. This is done by performing eigendecomposition on the covariance matrix of the point cloud coordinates, providing three eigenvectors and three eigenvalues. Our rotation then aligns the eigenvector corresponding to the highest eigenvalue to the y-axis basis vector (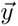 = [0, 1, 0]) and the eigenvector corresponding to the second highest highest eigenvalue to the x-axis basis vector (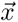 = [1, 0, 0]). As the covariance matrix is square and symmetric, the eigenvectors are guaranteed to be orthogonal and as such by definition the final eigenvector is aligned with the final basis (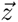 = [0, 0, 1]).

The second step taken is to fit a sphere to each population and subtract the centre of mass of this fitted sphere to provide a shared, central, coordinates origin for all dendrite populations regardless of neuropil. This works well, as both Medulla layer 10 and Lobula layer 1 are well approximated by a section on the surface of a sphere. As a result of this, the greatest extent of each populations points is aligned with the polar axis, and the y-axis, and the second with the equatorial axis, and the x-axis.

As a final step, we ensure that the anatomically dorsal population for both T4 and T5 is positioned within the northern hemisphere to ensure that both T4 and T5 dendrite populations are aligned with the same dorsal-ventral and anterior-posterior space. This final rotation is completed with reference to the column annotations within FAFB-FlyWire. This is done by identifying example dorsal, ventral, anterior and posterior columns as well as the T4 and T5 neurons associated with them and using these to ensure the correct global orientation. These available annotations assign neurons to a column in visual space, but do not include neuropil-specific spatial columns, necessitating this final step in order to facilitate the shared DV/AP alignment of the T4 and T5 dendrite populations.

These steps results in all T4 and T5 dendrites within both hemispheres being aligned so that the *y*-axis follows the Dorsal - Ventral axis and *x*-axis the Anterior-Posterior.

### Individual Dendrite Alignment and Scaling

We utilise a similar procedure to align and scale individual dendrites where applicable. When aligning individual dendrites, we calculate the covariance matrix using a robust M-estimation procedure based on a Huber-type influence function. This is done in order to minimise the influence of node outliers on eigenvectors as a single long outreaching sections may impact the eigenvector orientation within single dendrites. After Eigendecomposition of the covariance matrix, eigenvectors are aligned similarly to the global procedure so the primary eigen-axis aligns with the y-axis, the secondary eigen-axis to the *x*-axis and the third to the *z*-axis. Before calculating angles of rotation we ensure that each eigenvector is oriented along it’s positive target basis vector, in order to minimise rotation.

In order to scale point node positions we scale coordinates along their eigenvectors by *s*. Within Fig 2A, these eignevectors are the basis vectors in euclidean space due to the above described alignment. Again for Fig 2A, *s* is defined as:

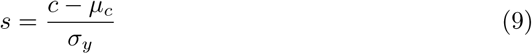

where *c* is the node coordinates of a single dendrite, *µ_c_* is the mean coordinate, and *σ_y_* is the mean standard deviation along the y-axis of the current population. This scales all dendrites along each axis so that 1 = *σ_y_*, placing the coordinates in standard deviation units. Alternatively, for unit variance scaling (Fig 7B, 7D and 7F) *s* is a vector equal to 1*/*var[*e_i_*] where var[*e_i_*] is the variance along each eigenvector, *e*. This scales dendrite to have equal variance along each axis.

### Branching Structure and Symmetry Quantification

Considering a tree graph with no nodes where deg^+^ = deg*^−^* = 1, as is the case for the reduced neuron tree-graph representations used here, let *n* = *|V |* and *L* be the number of leaves. Internal node number *I* = *n − L*. As (i) a *star* tree has one branch point and *L* = *n −* 1, and (ii) a *full binary* tree satisfies *L* = *I* + 1, hence *I* = (*n −* 1)*/*2, thus for any tree graph we have an upper bound:

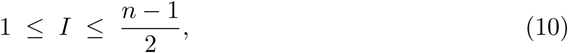

This provides limits between maximally condensed and maximally branching tree-graphs, and no tree-graph can exist outside of these bounds. The closer *I* is to (*n −* 1)*/*2, the closer to a full binary tree.

Tree asymmetry for a single node, *A_v_*, as defined in [26, 27], is extended similarly to [28] in order to weight the metric by the weight of the subtree defined at *v* with *k* ≥ 2 child nodes. In the case of deg^+^(*v*) > 2, we use a mean of all child pairs (*p, q*):

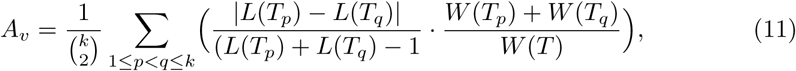

*A_v_* lies in [0, 1], where 0 indicates full symmetry in the branching structure.

### Surface Fitting, Volume, and Depth Measurement

In order to find the spanning volume of single dendrites, a convex hull is fit to the node coordinates of each dendrite using scipy.spatial.ConvexHull [73]. We reconstruct surfaces meshes of Medulla layer 10 and Lobula layer 1 via voxelization and marching cubes of all node coordinates in T4 (Medulla layer 10) or T5 (Lobula layer 1) dendrites. Step by step, we first compute the Axis-aligned bounding box for the given point cloud, extended by a fixed padding factor (2*µm*) to ensure a spatial margin. This is then discretized into 3*µm*^3^ voxels and each point is assigned to a voxel. 1 iteration of morphological dilation is applied in order to fill small gaps in the set of occupied voxels, and then unoccupied voxels are removed, retaining only the largest connected region of voxels. This binary grid is then converted to a continuous surface using the marching cubes algorithm as implemented in scikit-image. The resulting mesh is then cleaned by removing unreferenced nodes, duplicated or degenerate faces, filling holes in the mesh surface, and correcting face normals to ensure they are outwardly oriented. Surface meshes are smoothed using the Windowed Sinc method and a band pass filter with a width of 0.01*µm*, as implemented in [74].

In order to calculate the depth of any given point within a neuropil layer mesh, we first define two subsets of faces on the surface mesh which represent the two sides of the neuropil we are measuring depth across. We can define these sub-surfaces base on the orientation of face normals in relation to the shared coordinate origin. This is done by taking the dot product of each face normal with a vector from the coordinate origin. Faces with values ≥ 0.8 are treated as belonging to the outer mesh surface, and values ≤ −0.8 as the inner surface. A KDTree is calculated for both the inner and outer surface. For any given point, the nearest neighbour distance is calculated to both surfaces. We then calculate a normalized distance for the point from the inner surface. As T5 dendrites innervate Lobula layer 1 from the opposite side of curvature in comparison to T4 dendrites innervating Medulla layer 10, we invert T5 depths. Depth is calculated as follows:

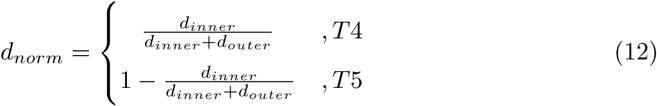

When generating neuropil surface meshes, a dilation step which fills in missing voxels is required in order ensure a single continuous surface is recovered. Due to the curvature of these neuropils, this results in a degree of padding within the concave region which is empty of dendrites. The extent of this padding differs between the two hemispheres, resulting in an offset when estimating neuropil layer depth. As such, we treat analysis of depth in the two hemispheres separately.

### Dendrite Root Point Nearest Neighbour Matching

In order to identify dendrite root point nearest neighbours we first assign subtype specific nearest neighbours using KDTrees within each type - subtype combination, allowing us to identify the distance between each T4a, for example, and the closest T4a (not including itself), T4b, T4c, and T4d dendrite root.

In order to determine the probability of a nearest neighbour being of either a specific classification (”opposite”, “same”, or “orthogonal”), or specific subtype, we calculate an optimal pairing within all T4 and all T5 dendrite root positions using a maximum-cardinality matching approach as implemented within the graph-tool python toolbox [75]. This method aims to match as many points as possible, maximising the number of pairs while simultaneously minimising the total euclidean distance between the matched pairs.

In order to account for depth as a possible confound within this procedure we project each set of points to the surface of their fitted sphere prior to calculating nearest neighbour distances as described above (S3 Fig).

### Geometry Analysis

When calculating angles between vectors in ℝ^3^ we project vectors [*v*_1_*, v*_2_] to the plane orthogonal to the normal vector, *n*, by taking the rejection of each vector from *n*:

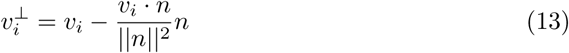

We then calculate the signed angle, *θ*, between *v*_1_ and *v*_2_ in the plane as:

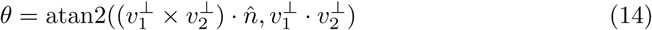

Where n̂ is the unit normal of the basis vector, 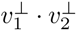 is the cosine of the angle. 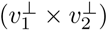· n̂ gives the signed sine, where the sign is determined by the right hand rule with respect to *n*. atan2(*y, x*) returns the signed angle *θ* ∈ (*−π, π*].

As defined in [76], the circular mean is calculated as:

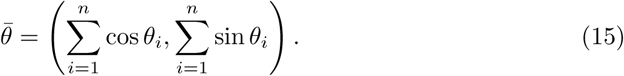

and Variance as:

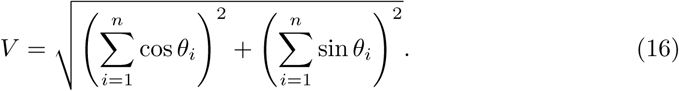

The Dihedral angle *β*, is the “the angle between the daughters’ half-plane and the plane formed by the parent segment and the line perpendicular to daughters’ bisection through the bifurcation point in the daughters’ plane” [22] (Fig 5L, insert). In order to calculate this for a set of parent and two child vectors [*p, c*_1_*, c*_2_] centred on their branching point, we first find the bisector, *b*, of *c*_1_ and *c*_2_ as:

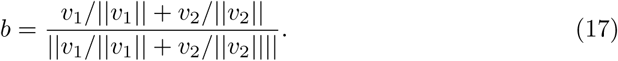

We then calculate the angle between *b* and *p* from the perspective of the plane defined by the normal vector, *n*, given by:

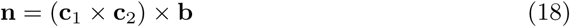

This results in the Dihedral angle *β* which is between 0 and *π*. At both extremes, this value represents bifurcations sitting on the same plane, however at 0 child vectors are on the plane oriented in the same direction as the parent relative to the bifurcation points, and at *π* they will be oriented away.

In order to generate bifurcation random null-models we randomly generate 1, 000, 000 sets of three 3-dimensional unit vectors. We ensure that one vector is oriented towards the negative *x*-axis to represent the parent section, and the other two are oriented along the positive *x*-axis. This ensures an approximation of the outward facing branching structure observed. Second, sets of bifurcation vectors are then scaled along their *z*-axis separately for T4 and T5 by the cube root of the mean variance explained by the third principle component, in order to mimic the natural flatness of the occupied space of T4 and T5 dendrites.

### Explorative Distribution Fitting

We divide dendrite populations by type (T4, T5), subtype (a,b,c,d) and section type (internal, external). We then fit using maximum likelihood, for each group, six test distributions in order to compare goodness of fit. The distributions used are as follows:

Wald (Inverse Gaussian) Distribution, with mean *µ* > 0 and shape *λ* > 0:

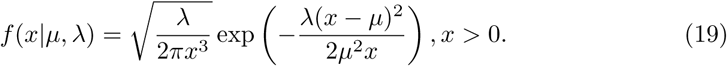

Log-normal Distribution, with log-mean *µ* ∈ ℝ and log-standard deviation *σ* > 0:

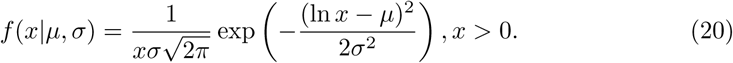

Gamma Distribution, with shape *k* > 0 and scale *θ* > 0:

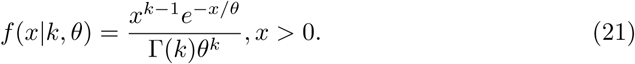

where Γ is the gamma function.

Exponential Distribution (Gamma distribution special case with *k* = 1) with scale *θ* > 0:

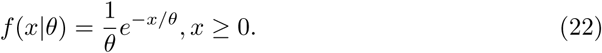

Log-logistic Distribution with shape *c* > 0 and scale *s* > 0:

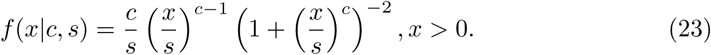

Minimum Weibull Distribution with shape *c* > 0 and scale *λ* > 0:

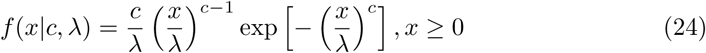

For each fitted distribution we calculate the Bayesian Information Criterion (BIC) as:

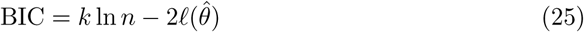

where *k* is the number of free parameters (2 for Wald, log-normal, gamma, log-logistic, and Weibull Distributions; 1 for exponential). *n* is the number of observations within the sample and *ℓ*(*θ̂*) is the maximised log-likelihood. Lower BIC indicates a better trade off between model fit and parsimony.

### Statistical Analysis

We test for hemisphere (Right, Left), neuron type (T4, T5) and subtype (a,b,c, and d) effects, as well as all interactions, on different measured variables using ordinary least squares (OLS) regression and analysis of variance (ANOVA) / covariance (ANCOVA). Models are fit with a full factorial structure, and sum of squares were evaluated using Type III tests. We additionally, for each effect (main effects and interaction terms), calculate the partial eta-squared (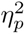) as an effect size, which quantifies the proportion of variance uniquely attributed to each factor given the model structure [18]:

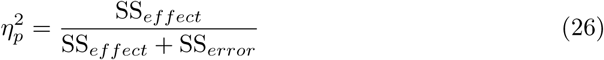

where SS is the sum of squares. We additionally report slopes (*β*) obtained from the regression model fit for meaningfully large main effects, as well as group specific slopes and their differences (Δ*β*) where applicable.

We additionally obtain, for each effect size, a non-parametric confidence interval obtained through stratified bootstrap resampling (1000 samples). *Post hoc* pairwise comparisons were conducted on marginal means, both at the main effect level and within interactions, and Cohen’s d was calculated for effect sizes:

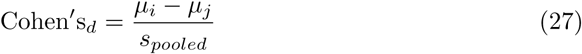

with

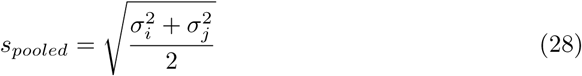

where *µ* is the group mean and *σ* the group standard deviation. Again, non-parametric confidence intervals using 1000 bootstrap resamples are calculated.

### Plotting Conventions

We make liberal use of the empirical probability mass functions (epmf) for plotting distributions throughout, without smoothing. To allow for comparison within plots, binning along the *x*-axis is the same for all epmf’s shown within a plot. Bin counts and ranges are adjusted manually. Error bars are presented in two forms, as specified in a plot by plot basis. In cases where a single observation is available per individual dendrite, a bootstrapped confidence interval around the mean in each bin is shown (1000 bootstrap replicates). Alternatively, when a repeated measure is available within each dendrite, a single epmf is generated for each individual dendrite and, in order to give an idea of individual variability across the population, error bars within each bin ar calculated as the asymmetric median absolute deviation (MAD*_a_*):

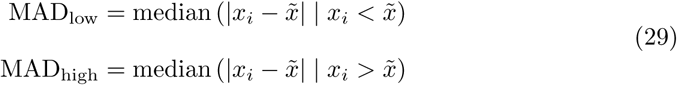

where *x̃* is the median within the respective bin. This illustrate variance in each bin across neurons. An Asymmetric MAD allows for the possibility that individual variability within each bin is skewed, and can be shown within the plot.

## Data and Code Availability

The open source Python toolboxes “fafbseg” [77], “skeletor” [69], and “navis” [68] were used to obtain and skeletonize neuron meshes. The code repository associated with this paper is available online (https://doi.org/10.5281/zenodo.21876360). This includes example Jupyter notebooks illustrating how to access the data from FAFB-FlyWire, download meshes, and skeletonize neurons using fafbseg and navis. Additionally, we provide the code required for going from an swc file representation of a complete neuron to individual extracted dendrites and code for the calculation of all metrics, as well as alignment and scaling steps implemented within the paper. This requires the “NeuRosetta” toolbox, available online (https://github.com/NikDrummond/NeuRosetta). This toolbox also includes the GUI used for manual verification of dendrites and dendrite annotation. Finally, we provide additional notebooks, as well as custom python code illustrating the implementation of analysis and plotting using standard python libraries. An additional datastore is available online (https://doi.org/10.5281/zenodo.21876510) which includes data files for all extracted dendrite metrics used within this manuscript.

## Supporting information

**S1 Fig.** Spanned Region of T4 and T5 Dendrites Separated by Hemispheres.

**S2 Fig.** Distribution and Correlations of Size Based Dendrite Metrics.

**S3 Fig.** Mi1 defined Medulla Columns within Layer 10.

**S4 Fig** Dendrite root nearest neighbour relationships for sphere projected dendrite root point nearest neighbours.

**S5 Fig** Subtype specific Bayesian Information Criterion (BIC) for all Fitted Distributions for Raw and Scaled Dendrite Section Cable Lengths.

**S1 Text.** Statistical analysis for S2 Fig, and Materials and Methods describing Mi1 based column mapping and spherical projections for dendrite root nearest neighbour analysis within S4 Fig. Table A includes counts of all available and included neuron morphologies before and after dendrite annotation, as well as counts of Mi1 neurons used for column mapping within S3 Fig.

## Acknowledgments

We are grateful to all members of the Borst Department for helpful comments and discussion of the manuscript and analysis. We thank the Princeton FlyWire team and members of the Murthy and Seung labs for development and maintenance of FlyWire (supported by BRAIN Initiative grant MH117815 to Murthy and Seung). We also acknowledge members of the Princeton FlyWire team and the FlyWire consortium for neuron proofreading. We also wish to acknowledge Michael Reiser, for sharing an initial dataset of manually reconstructed T4 and T5 neurons, as well as those tracers in Janelia who contributed to this manual reconstruction effort, which provided an initial dataset prior to automated reconstructions from FlyWire.

## Author Contributions

**N. Drummond** Conceptualisation, Data Curation, Formal Analysis, Methodology, Software, Visualisation, Writing - Original draft preparation, Writing - Review and Editing. **A. Zhao** Conceptualisation, Methodology, Writing - Review and Editing. **A. Borst** Conceptualisation, Funding Acquisition, Supervision, Writing - Review and Editing.

## References

1. Borst A, Groschner LN. How Flies See Motion. Annual Review of Neuroscience. 2023;46(Volume 46, 2023):17–37. doi:10.1146/annurev-neuro-080422-111929.

2. Haag J, Arenz A, Serbe E, Gabbiani F, Borst A. Complementary Mechanisms Create Direction Selectivity in the Fly. eLife. 2016;5:e17421. doi:10.7554/eLife.17421.

3. Maisak MS, Haag J, Ammer G, Serbe E, Meier M, Leonhardt A, et al. A Directional Tuning Map of Drosophila Elementary Motion Detectors. Nature. 2013;500(7461):212–216. doi:10.1038/nature12320.

4. Shinomiya K, Huang G, Lu Z, Parag T, Xu CS, Aniceto R, et al. Comparisons between the ON- and OFF-edge Motion Pathways in the Drosophila Brain. eLife. 2019;8:e40025. doi:10.7554/eLife.40025.

5. Gruntman E, Romani S, Reiser MB. The Computation of Directional Selectivity in the Drosophila OFF Motion Pathway;8:e50706. doi:10.7554/eLife.50706.

6. Gruntman E, Romani S, Reiser MB. Simple Integration of Fast Excitation and Offset, Delayed Inhibition Computes Directional Selectivity in Drosophila;21(2):250–257. doi:10.1038/s41593-017-0046-4.

7. Pinto-Teixeira F, Koo C, Rossi AM, Neriec N, Bertet C, Li X, et al. Development of Concurrent Retinotopic Maps in the Fly Motion Detection Circuit. Cell. 2018;173(2):485–498.e11. doi:10.1016/j.cell.2018.02.053.

8. Hörmann N, Schilling T, Ali AH, Serbe E, Mayer C, Borst A, et al. A Combinatorial Code of Transcription Factors Specifies Subtypes of Visual Motion-Sensing Neurons in Drosophila. Development. 2020;147(9):dev186296. doi:10.1242/dev.186296.

9. Matsliah A, Yu Sc, Kruk K, Bland D, Burke AT, Gager J, et al. Neuronal Parts List and Wiring Diagram for a Visual System. Nature. 2024;634(8032):166–180. doi:10.1038/s41586-024-07981-1.

10. Zhao A, Gruntman E, Nern A, Iyer N, Rogers EM, Koskela S, et al. Eye Structure Shapes Neuron Function in Drosophila Motion Vision. Nature. 2025;646(8083):135–142. doi:10.1038/s41586-025-09276-5.

11. Henning M, Ramos-Traslosheros G, Gür B, Silies M. Populations of Local Direction–Selective Cells Encode Global Motion Patterns Generated by Self-Motion. Science Advances. 2022;8(3):eabi7112. doi:10.1126/sciadv.abi7112.

12. Apitz H, Salecker I. Spatio-Temporal Relays Control Layer Identity of Direction-Selective Neuron Subtypes in Drosophila;9:2295. doi:10.1038/s41467-018-04592-z.

13. Dorkenwald S, McKellar CE, Macrina T, Kemnitz N, Lee K, Lu R, et al. FlyWire: Online Community for Whole-Brain Connectomics. Nature Methods. 2022;19(1):119–128. doi:10.1038/s41592-021-01330-0.

14. Dorkenwald S, Matsliah A, Sterling AR, Schlegel P, Yu Sc, McKellar CE, et al. Neuronal Wiring Diagram of an Adult Brain. Nature. 2024;634(8032):124–138. doi:10.1038/s41586-024-07558-y.

15. Schlegel P, Yin Y, Bates AS, Dorkenwald S, Eichler K, Brooks P, et al. Whole-Brain Annotation and Multi-Connectome Cell Typing of Drosophila. Nature. 2024;634(8032):139–152. doi:10.1038/s41586-024-07686-5.

16. Berg S, Beckett IR, Costa M, Schlegel P, Januszewski M, Marin EC, et al.. Sexual Dimorphism in the Complete Connectome of the Drosophila Male Central Nervous System; 2025.

17. Sullivan GM, Feinn R. Using Effect Size—or Why the P Value Is Not Enough. Journal of Graduate Medical Education. 2012;4(3):279–282. doi:10.4300/JGME-D-12-00156.1.

18. Lakens D. Calculating and Reporting Effect Sizes to Facilitate Cumulative Science: A Practical Primer for t-Tests and ANOVAs. Frontiers in Psychology. 2013;4. doi:10.3389/fpsyg.2013.00863.

19. Cohen J. Statistical Power Analysis for the Behavioral Sciences. 2nd ed. New York: Routledge; 2013.

20. Takemura Sy, Nern A, Chklovskii DB, Scheffer LK, Rubin GM, Meinertzhagen IA. The Comprehensive Connectome of a Neural Substrate for ‘ON’ Motion Detection in Drosophila. eLife. 2017;6:e24394. doi:10.7554/eLife.24394.

21. Ouyang X, Sutradhar S, Trottier O, Shree S, Yu Q, Tu Y, et al. Neurons Exploit Stochastic Growth to Rapidly and Economically Build Dense Radially Oriented Dendritic Arbors. bioRxiv. 2025; p. 2025.02.24.639873. doi:10.1101/2025.02.24.639873.

22. van Pelt J, Uylings HBM. The Flatness of Bifurcations in 3D Dendritic Trees: An Optimal Design. Frontiers in Computational Neuroscience. 2012;5:54. doi:10.3389/fncom.2011.00054.

23. Uylings HBM, Kuypers K, Diamond MC, Veltman WAM. Effects of Differential Environments on Plasticity of Dendrites of Cortical Pyramidal Neurons in Adult Rats. Experimental Neurology. 1978;62(3):658–677. doi:10.1016/0014-4886(78)90276-5.

24. Uylings HBM, Kuypers K, Veltman WAM. Environmental Influences on the Neocortex in Later Life. In: Corner MA, Baker RE, Vandepoll NE, Swaab DF, Uylings HBM, editors. Progress in Brain Research. vol. 48 of Maturation of the Nervous System. Elsevier; 1978. p. 261–274.

25. Kim Y, Sinclair R, De Schutter E. Local Planar Dendritic Structure: A Uniquely Biological Phenomenon? BMC Neuroscience. 2009;10(1):P4. doi:10.1186/1471-2202-10-S1-P4.

26. Van Pelt J, Uylings HBM, Verwer RWH, Pentney RJ, Woldenberg MJ. Tree Asymmetry—A Sensitive and Practical Measure for Binary Topological Trees. Bulletin of Mathematical Biology. 1992;54(5):759–784. doi:10.1016/S0092-8240(05)80142-9.

27. Costa LdF, Zawadzki K, Miazaki M, Viana MP, Taraskin S. Unveiling the Neuromorphological Space. Frontiers in Computational Neuroscience. 2010;4. doi:10.3389/fncom.2010.00150.

28. Kassraian-Fard P, Pfeiffer M, Bauer R. A Generative Growth Model for Thalamocortical Axonal Branching in Primary Visual Cortex. PLOS Computational Biology. 2020;16(2):e1007315. doi:10.1371/journal.pcbi.1007315.

29. Van Pelt J, Verwer RWH. Growth Models (Including Terminal and Segmental Branching) for Topological Binary Trees. Bulletin of Mathematical Biology. 1985;47(3):323–336. doi:10.1007/BF02459919.

30. Kim J, Kwon N, Chang S, Kim KT, Lee D, Kim S, et al. Altered Branching Patterns of Purkinje Cells in Mouse Model for Cortical Development Disorder. Scientific Reports. 2011;1:122. doi:10.1038/srep00122.

31. Cuntz H, Forstner F, Borst A, Häusser M. One Rule to Grow Them All: A General Theory of Neuronal Branching and Its Practical Application. PLOS Computational Biology. 2010;6(8):e1000877. doi:10.1371/journal.pcbi.1000877.

32. Kawaguchi Y, Karube F, Kubota Y. Dendritic Branch Typing and Spine Expression Patterns in Cortical Nonpyramidal Cells. Cerebral Cortex. 2006;16(5):696–711. doi:10.1093/cercor/bhj015.

33. Nowakowski RS, Hayes NL, Egger MD. Competitive Interactions during Dendritic Growth: A Simple Stochastic Growth Algorithm. Brain Research. 1992;576(1):152–156. doi:10.1016/0006-8993(92)90622-g.

34. Lindsay RD, Scheibel AB. Quantitative Analysis of the Dendritic Branching Pattern of Small Pyramidal Cells from Adult Rat Somesthetic and Visual Cortex. Experimental Neurology. 1974;45(3):424–434. doi:10.1016/0014-4886(74)90149-6.

35. Rößler N, Jungenitz T, Sigler A, Bird A, Mittag M, Rhee JS, et al. Skewed Distribution of Spines Is Independent of Presynaptic Transmitter Release and Synaptic Plasticity, and Emerges Early during Adult Neurogenesis. Open Biology. 2023;13(8):230063. doi:10.1098/rsob.230063.

36. Loewenstein Y, Kuras A, Rumpel S. Multiplicative Dynamics Underlie the Emergence of the Log-Normal Distribution of Spine Sizes in the Neocortex In Vivo. Journal of Neuroscience. 2011;31(26):9481–9488. doi:10.1523/JNEUROSCI.6130-10.2011.

37. Diyali B, Kumar D, Singh S. Discriminating between Log-Normal and Log-Logistic Distributions in the Presence of Type-II Censoring. Computational Statistics. 2024;39(3):1459–1483. doi:10.1007/s00180-023-01351-7.

38. Gupta RD, Kundu D. Discriminating between Weibull and Generalized Exponential Distributions. Computational Statistics & Data Analysis. 2003;43(2):179–196. doi:10.1016/S0167-9473(02)00206-2.

39. de Berg M, van Kreveld M, Overmars M, Schwarzkopf O. Computational Geometry: Algorithms and Applications. Springer Science & Business Media; 2000.

40. Buffry AD, Currea JP, Franke-Gerth FA, Palavalli-Nettimi R, Bodey AJ, Rau C, et al. Evolution of Compound Eye Morphology Underlies Differences in Vision between Closely Related Drosophila Species. BMC Biology. 2024;22(1):67. doi:10.1186/s12915-024-01864-7.

41. Posnien N, Hopfen C, Hilbrant M, Ramos-Womack M, Murat S, Schönauer A, et al. Evolution of Eye Morphology and Rhodopsin Expression in the Drosophila Melanogaster Species Subgroup. PLOS ONE. 2012;7(5):e37346. doi:10.1371/journal.pone.0037346.

42. Choi BJ, Chen YC, Desplan C. Retinal Calcium Waves Coordinate Uniform Tissue Patterning of the Drosophila Eye. Science. 2025;390(6775):eady5541. doi:10.1126/science.ady5541.

43. Baltruschat L, Tavosanis G, Cuntz H. A Developmental Stretch-and-Fill Process That Optimises Dendritic Wiring; 2020. https://www.biorxiv.org/content/10.1101/2020.07.07.191064v1.

44. Sugimura K, Shimono K, Uemura T, Mochizuki A. Self-Organizing Mechanism for Development of Space-filling Neuronal Dendrites;3(11):e212. doi:10.1371/journal.pcbi.0030212.

45. Han X, Wang M, Liu C, Trush O, Takayama R, Akiyama T, et al. DWnt4 and DWnt10 Regulate Morphogenesis and Arrangement of Columnar Units via Fz2/PCP Signaling in the \mkbibemphDrosophila Brain. Cell Reports. 2020;33(4):108305. doi:10.1016/j.celrep.2020.108305.

46. Urbach R, Technau GM. Molecular Markers for Identified Neuroblasts in the Developing Brain of Drosophila. Development. 2003;130(16):3621–3637. doi:10.1242/dev.00533.

47. Lee YJ, Yang CP, Miyares RL, Huang YF, He Y, Ren Q, et al. Conservation and Divergence of Related Neuronal Lineages in the Drosophila Central Brain. eLife. 2020;9:e53518. doi:10.7554/eLife.53518.

48. Wolterhoff N, Hiesinger PR. Synaptic Promiscuity in Brain Development. Current Biology. 2024;34(3):R102–R116. doi:10.1016/j.cub.2023.12.037.

49. Frank SA. The Common Patterns of Nature. Journal of Evolutionary Biology. 2009;22(8):1563–1585. doi:10.1111/j.1420-9101.2009.01775.x.

50. Daley DJ, Vere-Jones D. An Introduction to the Theory of Point Processes. Vol. I. 2nd ed. Probability and Its Applications (New York). New York: Springer-Verlag; 2003.

51. van Veen MP, van Pelt J. Terminal and Intermediate Segment Lengths in Neuronal Trees with Finite Length. Bulletin of Mathematical Biology. 1993;55(2):277–294. doi:10.1007/BF02460884.

52. van Elburg RAJ. Stochastic Continuous Time Neurite Branching Models with Tree and Segment Dependent Rates. Journal of Theoretical Biology. 2011;276(1):159–173. doi:10.1016/j.jtbi.2011.01.039.

53. Chou ZZ, Yu GJ, Berger TW. Generation of Granule Cell Dendritic Morphologies by Estimating the Spatial Heterogeneity of Dendritic Branching. Frontiers in Computational Neuroscience. 2020;14. doi:10.3389/fncom.2020.00023.

54. Stürner T, Tatarnikova A, Mueller J, Schaffran B, Cuntz H, Zhang Y, et al. Transient Localization of the Arp2/3 Complex Initiates Neuronal Dendrite Branching in Vivo. Development. 2019;146(7):dev171397. doi:10.1242/dev.171397.

55. Hu X, Viesselmann C, Nam S, Merriam E, Dent EW. Activity-Dependent Dynamic Microtubule Invasion of Dendritic Spines. The Journal of Neuroscience: The Official Journal of the Society for Neuroscience. 2008;28(49):13094–13105. doi:10.1523/JNEUROSCI.3074-08.2008.

56. Dehmelt L, Smart FM, Ozer RS, Halpain S. The Role of Microtubule-Associated Protein 2c in the Reorganization of Microtubules and Lamellipodia during Neurite Initiation. The Journal of Neuroscience: The Official Journal of the Society for Neuroscience. 2003;23(29):9479–9490. doi:10.1523/JNEUROSCI.23-29-09479.2003.

57. Limpert E, Stahel WA, Abbt M. Log-Normal Distributions across the Sciences: Keys and Clues: On the Charms of Statistics, and How Mechanical Models Resembling Gambling Machines Offer a Link to a Handy Way to Characterize Log-Normal Distributions, Which Can Provide Deeper Insight into Variability and Probability—Normal or Log-Normal: That Is the Question. BioScience. 2001;51(5):341–352. doi:10.1641/0006-3568(2001)051[0341:LNDATS]2.0.CO;2.

58. Koch AL. The Logarithm in Biology 1. Mechanisms Generating the Log-Normal Distribution Exactly. Journal of Theoretical Biology. 1966;12(2):276–290. doi:10.1016/0022-5193(66)90119-6.

59. Statman A, Kaufman M, Minerbi A, Ziv NE, Brenner N. Synaptic Size Dynamics as an Effectively Stochastic Process. PLOS Computational Biology. 2014;10(10):e1003846. doi:10.1371/journal.pcbi.1003846.

60. Shomar A, Geyrhofer L, Ziv NE, Brenner N. Cooperative Stochastic Binding and Unbinding Explain Synaptic Size Dynamics and Statistics. PLOS Computational Biology. 2017;13(7):e1005668. doi:10.1371/journal.pcbi.1005668.

61. Shree S, Sutradhar S, Trottier O, Tu Y, Liang X, Howard J. Dynamic Instability of Dendrite Tips Generates the Highly Branched Morphologies of Sensory Neurons. Science Advances. 2022;8(26):eabn0080. doi:10.1126/sciadv.abn0080.

62. Nern A, Loesche F, Takemura Sy, Burnett LE, Dreher M, Gruntman E, et al.. Connectome-Driven Neural Inventory of a Complete Visual System; 2024. https://www.biorxiv.org/content/10.1101/2024.04.16.589741v2.

63. Zipursky SL, Sanes JR. Chemoaffinity Revisited: Dscams, Protocadherins, and Neural Circuit Assembly. Cell. 2010;143(3):343–353. doi:10.1016/j.cell.2010.10.009.

64. Jan YN, Jan LY. Branching out: Mechanisms of Dendritic Arborization. Nature Reviews Neuroscience. 2010;11(5):316–328. doi:10.1038/nrn2836.

65. Goodman CS, Shatz CJ. Developmental Mechanisms That Generate Precise Patterns of Neuronal Connectivity. Cell. 1993;72:77–98. doi:10.1016/S0092-8674(05)80030-3.

66. Zhao A, Nern A, Koskela S, Dreher M, Erginkaya M, Laughland CW, et al. A Comprehensive Neuroanatomical Survey of the Drosophila Lobula Plate Tangential Neurons with Predictions for Their Optic Flow Sensitivity. bioRxiv. 2023; p. 2023.10.16.562634. doi:10.1101/2023.10.16.562634.

67. Zheng Z, Lauritzen JS, Perlman E, Robinson CG, Nichols M, Milkie D, et al. A Complete Electron Microscopy Volume of the Brain of Adult Drosophila Melanogaster. Cell. 2018;174(3):730–743.e22. doi:10.1016/j.cell.2018.06.019.

68. Schlegel P, Barnes C, Loesche F, Champion A, dokato, Jagannathan S, et al.. Navis-Org/Navis: Version 1.10.0; 2025. Zenodo.

69. Schlegel P, Gokaslan A, Kazimiers T. Navis-Org/Skeletor: Version 1.3.0; 2024. Zenodo.

70. Meier M, Borst A. Extreme Compartmentalization in a Drosophila Amacrine Cell. Current biology: CB. 2019;29(9):1545–1550.e2. doi:10.1016/j.cub.2019.03.070.

71. Schneider-Mizell CM, Gerhard S, Longair M, Kazimiers T, Li F, Zwart MF, et al. Quantitative Neuroanatomy for Connectomics in Drosophila. eLife. 2016;5:e12059. doi:10.7554/eLife.12059.

72. Bates AS, Schlegel P, Roberts RJV, Drummond N, Tamimi IFM, Turnbull R, et al. Complete Connectomic Reconstruction of Olfactory Projection Neurons in the Fly Brain. Current Biology. 2020;30(16):3183–3199.e6. doi:10.1016/j.cub.2020.06.042.

73. Virtanen P, Gommers R, Oliphant TE, Haberland M, Reddy T, Cournapeau D, et al. SciPy 1.0: Fundamental Algorithms for Scientific Computing in Python. Nature Methods. 2020;17(3):261–272. doi:10.1038/s41592-019-0686-2.

74. Musy M, Jacquenot G, Dalmasso G, Lee J, Pujol L, Soltwedel J, et al.. marcomusy/vedo: v2025.5.4; 2025. Available from: 10.5281/zenodo.15630979.

75. Peixoto TP. Descriptive vs. Inferential Community Detection in Networks: Pitfalls, Myths and Half-Truths. Elements in the Structure and Dynamics of Complex Networks. 2023;doi:10.1017/9781009118897.

76. Jammalamadaka SR, SenGupta A. Topics in Circular Statistics. vol. 5. World Scientific Publishing Co. Pte. Ltd.; 2001.

77. Schlegel P, Perlman E. Navis-Org/Fafbseg-Py: Version 3.0.10; 2024. Zenodo.

