## Supplemental Text for "Population Morphology Implies a Common Developmental Blueprint for *Drosophila* Motion Detectors"

### Supplementary Material

#### Supplementary Figure Captions

**Fig 1.** Spanned region of T4 and T5 Dendrites Separated Hemispheres. a) T4 and T5 dendrites locally aligned and scaled, overlayed by subtype identically to Fig. 2,a, except split by left and right hemisphere.

**Fig 2.** Distribution and Correlations of Size Based Dendrite Metrics. a) Probability mass function of total dendrite cable lengths. b) Spatial distribution of total dendrite cable length. c) Correlation between total dendrite cable length and dendrite convex hull volume. d) Probability mass function of total number of dendrite sections. e) Spatial distribution of total number of dendrite sections. f) Correlation between total number of dendrite sections and convex hull volume.

**Fig 3.** Mi1 defined Medulla Columns within Layer 10. a) Example set of Mi1 surface meshes for $p = 6$ within the FlyWire column coordinate system. *Inset*: Single example Mi1 with fitted cylinder used for length and radius calculation. b) Mean distance to column neighbours within the left hemisphere. c) as in (b), for the right hemisphere. d) Length of fitted column cylinders from Mi1 neurons within the left hemisphere Medulla layer 10. e) as in (d), for the right hemisphere. f) Radius of fitted column cylinders from Mi1 neurons within the left hemisphere Medulla layer 10. g) As in (f), but for the right hemisphere Medulla layer 10.

**Fig 4.** Dendrite root nearest neighbour relationships for sphere projected dendrite root point nearest neighbours. a) Nearest neighbour distance of dendrite root points within T4 dendrites, using matching as in (Main Fig. 4a, left), grouped into horizontal (a, b subtypes, solid lines) and vertical (c, d subtypes, dashed lines). "Opposite" distributions denote opposingly oriented subtypes (eg, a→b). "Same" subtypes denote same subtype combinations (eg, a→a). "Orthogonal" denotes orthogonally oriented subtypes (eg, a→c and a→d). b) Probability of the nearest neighbour being of each of the "Opposite", "Same", and "Orthogonal", inferred using nearest neighbour matching derived from (Main Fig. 4a, right) within T4 dendrites. Dashed lines denote approximate chance probability for each group (0.25 for "Opposite" and "Same" groups, 0.5 for "Orthogonal" groups). c) Subtype specific nearest neighbour probabilities inferred from nearest neighbour matching derived from (Main Fig. 4a, right) within T4 dendrites. d-f) as in a-c but for T5 dendrites.

**Fig 5.** Subtype specific Bayesian Information Criterion (BIC) for all Fitted Distributions for Raw and Scaled Dendrite Section Cable Lengths. a) Raw internal section length BIC scores across subtypes for all fitted distributions. Fitted distribution names are the same as in (c) along the x-axis. b) Variance scaled internal section length BIC scores across al subtypes for al fitted distributions. Fitted distribution names are the same as in (d) along the x-axis. c) as in (a) but for external (terminal) section cable length. d) as in (b) for external (terminal) section cable length. Within all plots, the gray point denote the mean BIC score across subtypes. Error bars show ±1 standard deviation from the mean BIC. The dashed gray line shows the mean BIC for the 'winning' fitted distribution, that with the lowest value across all fitted distribution, Gamma for internal sections and log-normal for external sections.

#### Supplementary Results

##### Statistical Analysis of Dendrite Size Metrics

When comparing total dendrite cable length, no meaningfully large statistical differences are observed between hemispheres, types, or subtypes ($\eta_p^2 < 0.01$, Supplementary Fig 2A). As noted when considering dendrite convex hull volumes, the distribution of T4 dendrite total cable length shows the same asymmetry. Additionally dendrites with greater cable lengths are concentrated within the anterior-ventral region of the Medulla (Supplementary Fig 2B). When total dendrite cable length is used as a covariate predictor for dendrite convex hull volume (Supplementary Fig 2C) no meaningfully large statistical differences are found between hemisphere, neuron type, or subtype ($\eta_p^2 < 0.01$). However, total cable length is strong predictor of total dendrite convex hull volume, with a large effect size ($\eta_p^2 = 0.271, \mathrm{CI} : [0.226, 0.330]$). With a coefficient of $\beta = 2.773$ this indicates that, on average, a single dendrite fills $1\mu m^3$ with $\sim 2.773\mu m$ of dendritic cable.

When considering the number of sections within a dendrite, edges between the dendrite root point, branch points, and terminal points, we find no meaningfully large differences between hemispheres, neuron type or subtype ($\eta_p^2, 0.01$, Fig 2D). Again, as in dendrite convex hull volume, and dendrite total cable length, we observe a skewed distribution within T4 not present in T5, which is concentrated within the anterior-ventral region of the Medulla (Fig 2E). When considering dendrite section count as a covariate predictor of dendrite convex hull volume, we again find no meaningfully large effects of hemisphere, neuron type, or subtype, but do find that section count is a strong predictor of dendrite convex hull volume (Fig 2F), with a large effect size ($\eta_p^2 = 0.191, \mathrm{CI} : [0.152, 0.231]$). With a coefficient of $\beta = 3.689$ this indicates that each $\mu m^3$ of dendrite spanned volume, a dendrite has $\sim 3.689$ within, however the interpretation of this slope is not so intuitively interpretable as in the convex hull volume and dendrite cable length relationship.

#### Supplementary Methods

##### Column Mapping

We use Mi1 neurons in order to identify columns within the right and left Medulla. Using the surface mesh of each Mi1, we first subset the mesh to the points within the relevant surface layer mesh for each hemisphere. This provides a point cloud of each Mi1 neuron within Medulla layer 10 (Fig 3A).

For each Mi1 point cloud, similar to dendrite alignment steps taken, the internal axis are determined using principal component analysis, and the first principal component is used as the primary internal axis. Column length was estimated from the 99th percentile of the absolute coordinate projections onto this axis, while column radius was estimated from the 95th percentile of the perpendicular distances from the axis. percentile based estimates were used in order to reduce the influence of outliers within each point cloud.

In order to define the mean distance to neighbouring column, we rely on the Mi1 column annotations available from FlyWire, available at `https://https://codex.flywire.ai/app/visual_columns_map?dataset=fafb`. This column map uses a $[p, q]$ hexagonal coordinate system, which we transform to cubic $[q, r, s]$ coordinates:

$$\begin{pmatrix} q \\ r \\ s \end{pmatrix} = \begin{pmatrix} 1 & -1 \\ -1 & 0 \\ 0 & 1 \end{pmatrix} \begin{pmatrix} p \\ q \end{pmatrix} \tag{1}$$

Given the cubic coordinates of a single column within the hexagonal grid, we use the six nearest neighbour displacement vectors to the cubic coordinates of each of its neighbours: $(0, -1, 1), (1, -1, 0), (1, 0, -1), (0, 1, -1), (-1, 1, 0), (-1, 0, 1)$. For each coordinate $c = (q, r, s)$, neighbour coordinates are then obtained as $c + d$, where $d$ is one of the six displacement vectors. Given these neighbour coordinates we can then find the column which matches these coordinates, excluding neighbour columns which do not have an allocated Mi1 in FlyWire.

Given then a column and each of its existing neighbours, we use the centre of each Mi1 point cloud, and calculate the mean distance between the central column and each of its existing neighbours. These analysis are performed within each hemisphere separately.

##### Spherical Projection

In order to assess the impact of neuropil layer depth on dendrite root point nearest neighbour analysis we project dendrite root points onto a spherical surface. We re-use the spheres fitted to neuropil and hemisphere specific dendrites used for global dendrite alignment. given the dendrite root point $x$, we transform it by:

$$x' = c + R\frac{x - c}{||x - c||} \tag{2}$$

Where $R$ is the radius of the fitted sphere, and $c$ the centre, which in our case is the coordinate origin, and as such is 0. This maps all dendrite root points onto a sphere of radius $R$,c entered at $c$.

#### Data and Code Availability

Mi1 neuron IDs and column annotations are available from FlyWire (`https://codex.flywire.ai/?dataset=fafb`). Statistical analysis and plotting was carried out in the same manner as within the main paper, and code is available in the same associated GitHub repository. Custom python code for column maps is also available within this repository. Hexagonal heatmaps, and cubic coordinate transforms are implemented using the "HexCraft" toolbox, available at `https://github.com/NikDrummond/HexCraft`

#### Supplementary Table A

| Hemisphere | Subtype | Count in Flywire | Count Used | Proportion Used |
|---|---|---|---|---|
| R | T4a | 737 | 715 | 0.97 |
| R | T4b | 748 | 722 | 0.97 |
| R | T4c | 842 | 825 | 0.98 |
| R | T4d | 777 | 764 | 0.98 |
| R | T5a | 743 | 706 | 0.95 |
| R | T5b | 760 | 728 | 0.96 |
| R | T5c | 766 | 734 | 0.96 |
| R | T5d | 727 | 711 | 0.98 |
| L | T4a | 719 | 697 | 0.97 |
| L | T4b | 758 | 727 | 0.96 |
| L | T4c | 869 | 846 | 0.98 |
| L | T4d | 794 | 781 | 0.98 |
| L | T5a | 740 | 709 | 0.96 |
| L | T5b | 754 | 724 | 0.96 |
| L | T5c | 770 | 730 | 0.95 |
| L | T5d | 742 | 719 | 0.97 |
| R | Mi1 | 796 | na | na |
| L | Mi1 | 785 | na | na |

**Table A.** Counts of T4 and T5 dendrites by subtype, and Mi1 neurons used within the study and available in flywire. na with Mi1 counts indicates that all are used.
