## Supplemental Figures for "Population Morphology Implies a Common Developmental Blueprint for *Drosophila* Motion Detectors"

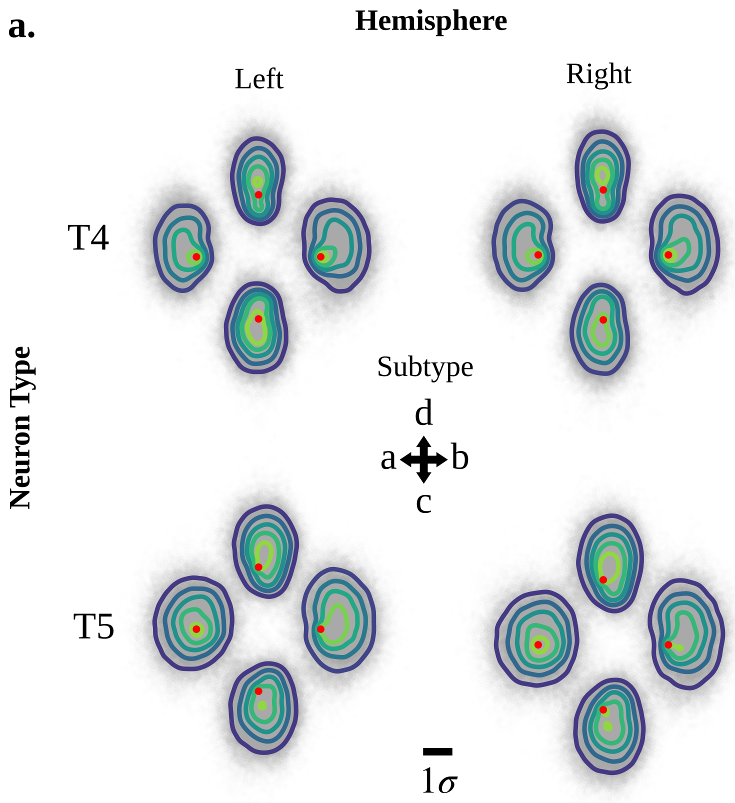


**Fig 1.** Spanned region of T4 and T5 Dendrites Separated Hemispheres. a) T4 and T5 dendrites locally aligned and scaled, overlayed by subtype identically to Fig. 2,a, except split by left and right hemisphere.

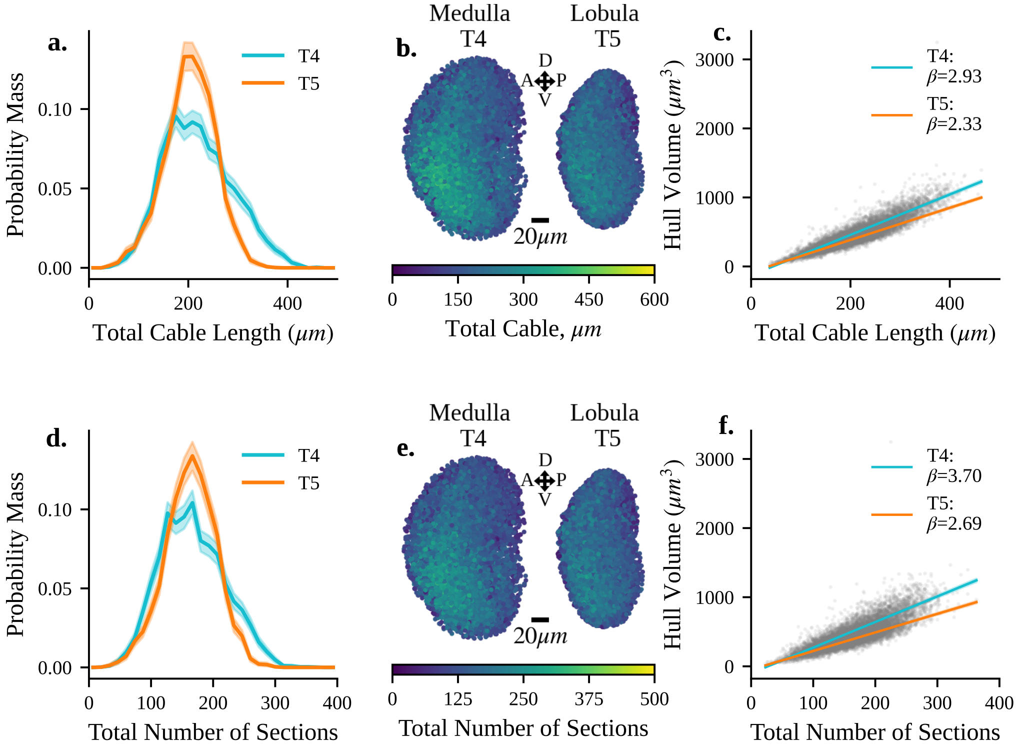


**Fig 2.** Distribution and Correlations of Size Based Dendrite Metrics. a) Probability mass function of total dendrite cable lengths. b) Spatial distribution of total dendrite cable length. c) Correlation between total dendrite cable length and dendrite convex hull volume. d) Probability mass function of total number of dendrite sections. e) Spatial distribution of total number of dendrite sections. f) Correlation between total number of dendrite sections and convex hull volume.

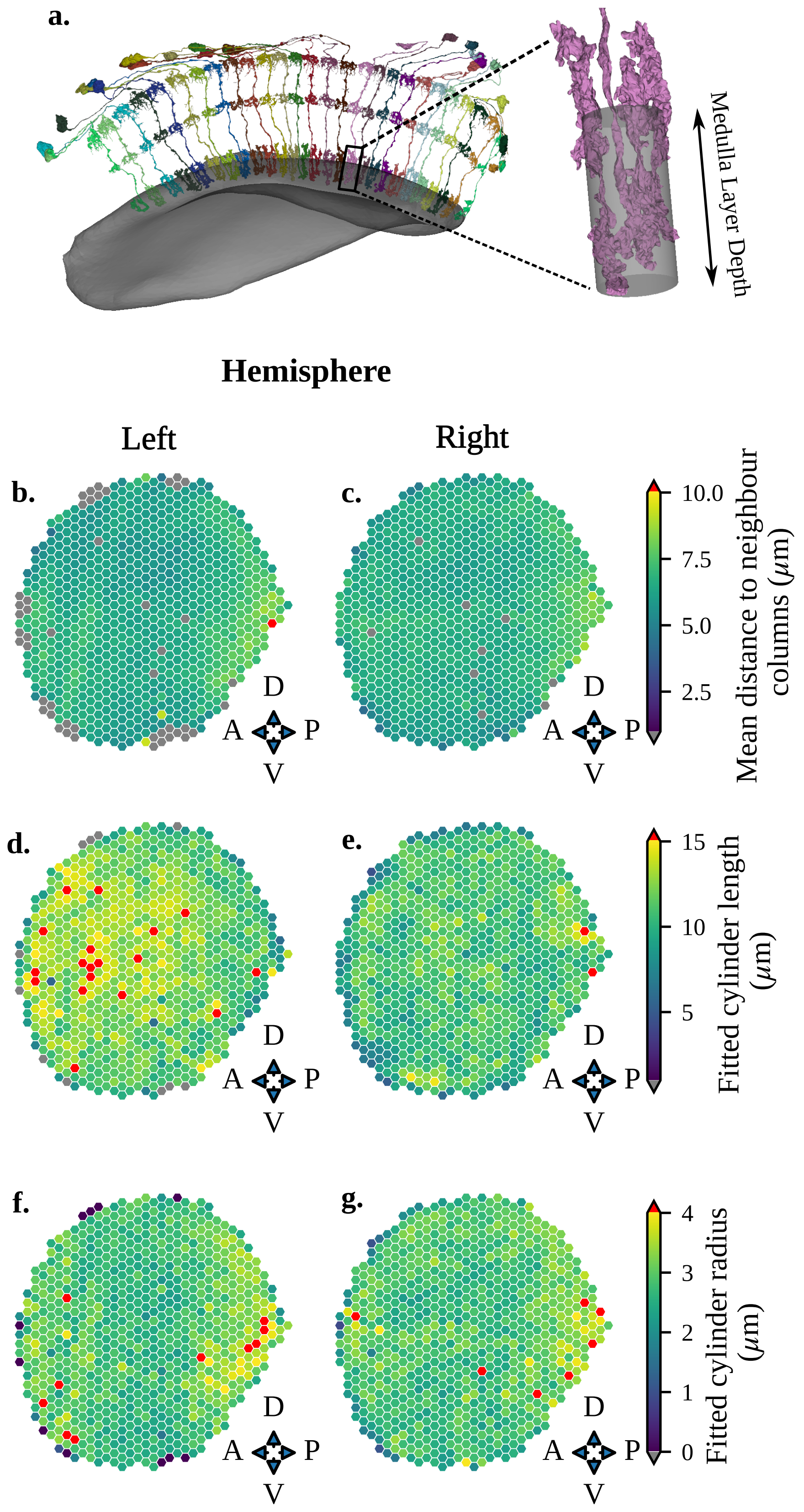
a.
Medulla Layer Depth
Hemisphere
Left
Right
b.
c.
d.
e.
f.
g.
D
A
P
V
Mean distance to neighbour columns ($\mu m$)
10.0
7.5
5.0
2.5
Fitted cylinder length ($\mu m$)
15
10
5
Fitted cylinder radius ($\mu m$)
4
3
2
1
0

**Fig 3.** Mi1 defined Medulla Columns within Layer 10. a) Example set of Mi1 surface meshes for $p = 6$ within the FlyWire column coordinate system. *Inset*: Single example Mi1 with fitted cylinder used for length and radius calculation. b) Mean distance to column neighbours within the left hemisphere. c) as in (b), for the right hemisphere. d) Length of fitted column cylinders from Mi1 neurons within the left hemisphere Medulla layer 10. e) as in (d), for the right hemisphere. f) Radius of fitted column cylinders from Mi1 neurons within the left hemisphere Medulla layer 10. g) As in (f), but for the right hemisphere Medulla layer 10.

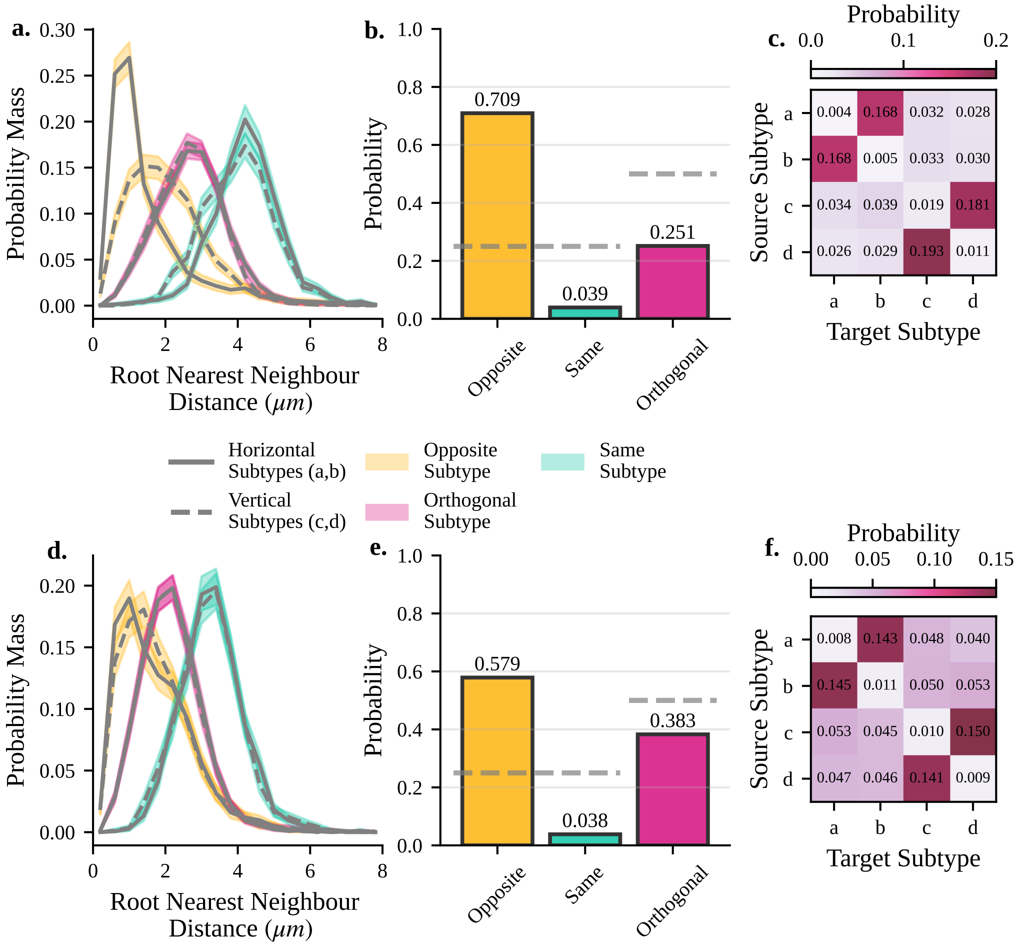


**Fig 4.** Dendrite root nearest neighbour relationships for sphere projected dendrite root point nearest neighbours. a) Nearest neighbour distance of dendrite root points within T4 dendrites, using matching as in (Main Fig. 4a, left), grouped into horizontal (a, b subtypes, solid lines) and vertical (c, d subtypes, dashed lines). ”Opposite” distributions denote opposingly oriented subtypes (eg, a→b). ”Same” subtypes denote same subtype combinations (eg, a→a). ”Orthogonal” denotes orthogonally oriented subtypes (eg, a→c and a→d). b) Probability of the nearest neighbour being of each of the ”Opposite”, ”Same”, and ”Orthogonal”, inferred using nearest neighbour matching derived from (Main Fig. 4a, right) within T4 dendrites. Dashed lines denote approximate chance probability for each group (0.25 for ”Opposite” and ”Same” groups, 0.5 for ”Orthogonal” groups). c) Subtype specific nearest neighbour probabilities inferred from nearest neighbour matching derived from (Main Fig. 4a, right) within T4 dendrites. d-f) as in a-c but for T5 dendrites.

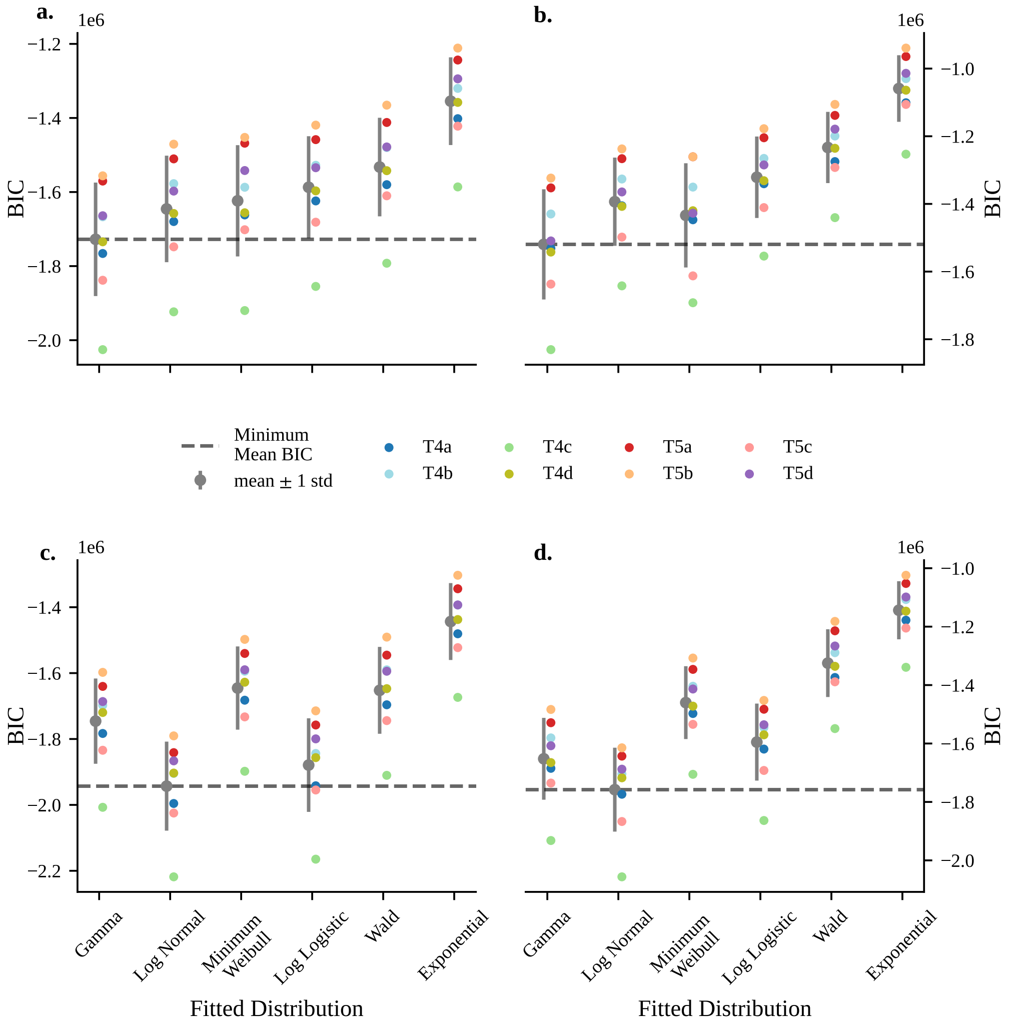


**Fig 5.** Subtype specific Bayesian Information Criterion (BIC) for all Fitted Distributions for Raw and Scaled Dendrite Section Cable Lengths. a) Raw internal section length BIC scores across subtypes for all fitted distributions. Fitted distribution names are the same as in (c) along the x-axis. b) Variance scaled internal section length BIC scores across al subtypes for al fitted distributions. Fitted distribution names are the same as in (d) along the x-axis. c) as in (a) but for external (terminal) section cable length. d) as in (b) for external (terminal) section cable length. Within all plots, the gray point denote the mean BIC score across subtypes. Error bars show ±1 standard deviation from the mean BIC. The dashed gray line shows the mean BIC for the 'winning' fitted distribution, that with the lowest value across all fitted distribution, Gamma for internal sections and log-normal for external sections.
